# The extracellular chaperone Clusterin enhances Tau aggregate seeding in a cellular model

**DOI:** 10.1101/2021.07.16.452659

**Authors:** Patricia Yuste-Checa, Victoria A. Trinkaus, Irene Riera-Tur, Rahmi Imamoglu, Theresa Schaller, Huping Wang, Irina Dudanova, Mark S. Hipp, Andreas Bracher, F. Ulrich Hartl

**Author notes:** Present address: Institute for Molecular Medicine, University Medical Center of the Johannes Gutenberg-University Mainz, Mainz, Germany (Theresa Schaller). Present address: Department of Biomedical Sciences of Cells and Systems, University Medical Center Groningen, University of Groningen, Groningen, The Netherlands (Mark S. Hipp). Present address: School of Medicine and Health Sciences, Carl von Ossietzky University Oldenburg, Oldenburg, Germany (Mark S. Hipp).

## Abstract

Spreading of aggregate pathology across brain regions acts as a driver of disease progression in Tau-related neurodegeneration, including Alzheimer’s disease (AD) and frontotemporal dementia. Aggregate seeds released from affected cells are internalized by naïve cells and induce the prion-like templating of soluble Tau into neurotoxic aggregates. Here we show in a cellular model system and in neurons that Clusterin, an abundant extracellular chaperone, strongly enhances Tau aggregate seeding. Upon interaction with Tau aggregates, Clusterin stabilizes highly potent, soluble seed species. Tau/Clusterin complexes enter recipient cells via endocytosis and compromise the endolysosomal compartment, allowing transfer to the cytosol where they propagate aggregation of endogenous Tau. Thus, upregulation of Clusterin, as observed in AD patients, may enhance Tau seeding and possibly accelerate the spreading of Tau pathology.

## Introduction

Progression of several neurodegenerative diseases, prominently including tauopathies such as frontotemporal dementia (FTD) and Alzheimer’s disease (AD), is driven by spreading of aggregate pathology across brain regions in a prion-like seeding mechanism^1–5^. Tau aggregate spreading involves the exposure of aggregate seeds to the extracellular milieu^6, 7^, suggesting that extracellular protein quality control factors may modulate disease progression^8^. Clusterin (Clu; apolipoprotein J) is a ~70 kDa glycoprotein with chaperone-like properties found abundantly in plasma and extracellular fluid^8–10^. During its passage through the secretory pathway, immature Clu is extensively N-glycosylated and cleaved into α and β-chains, which remain linked by disulfide bonds (Supplementary Fig. 1a). Although mainly a secreted chaperone, Clu is also found intracellularly under stress conditions^9^. Clu stabilizes unfolded proteins against aggregation and can inhibit fibril formation of amyloid β (Aβ) and other amyloidogenic proteins *in vitro*, consistent with the function of an ATP-independent ‘holdase’ chaperone^10–15^.

The *CLU* gene ranks third among genetic risk factors for late onset AD^16, 17^. However, the mechanism by which Clu modulates AD pathology remains unclear, as Clu has been associated with both neuroprotective and neurotoxic effects in AD^15, 18–25^. Clu protein expression is upregulated in AD patient brain and cerebrospinal fluid^26, 27^, localizing with Aβ deposits in senile plaques^28–30^. Evidence has been presented that Clu can mediate uptake of Aβ via the endosomal pathway by microglia and is involved in clearance of Aβ via the blood-brain barrier^31–33^. On the other hand, enhancement of Aβ toxicity by Clu has also been reported^24, 25^ and elevated plasma levels of Clu were found to be associated with rapid progression of AD, suggesting that Clu could be a driver of pathology^34, 35^. Little is known about a possible role of Clu in the progression of Tau pathology, which strongly correlates with the severity of AD^36–38^. Interestingly, Clu was identified as an interactor of soluble Tau in AD brain^39^. More recently, it was shown that Clu also colocalizes with intracellular Tau aggregates and may provide a protective function by inhibiting fibril formation^40^. Given the predominant role of extracellular Clu as the chaperone active form, it remains to be understood whether Clu modulates transcellular Tau seeding and influences overall pathology.

Here we analyzed the effect of Clu on the seeding competence of Tau aggregates formed *in vitro* and in cells. Our results show that Clu can strongly enhance Tau aggregate propagation by binding and stabilizing seeding active Tau species for cellular uptake. Thus, upregulation of Clu in AD has the potential to accelerate disease progression by enhancing the seeding competence of Tau aggregates.

## Results

### Clusterin potentiates seeding of Tau aggregates

To test whether Clu interferes with aggregate seeding of Tau, we purified chaperone-active Clu upon recombinant expression in HEK293-EBNA cells^10, 14^ (Supplementary Fig. 1b, c). We measured aggregate seeding with TauRD-YT cells, a HEK293T cell line stably co-expressing the repeat domain of Tau (TauRD; residues 244-372 with FTD mutations P301L/V337M) fused to YFP or mTuquoise2, whose co-aggregation during fibril formation results in fluorescence resonance energy transfer (FRET)^41^ (Fig. 1a). Seed aggregates were generated with recombinant, cysteine-free TauRD (Tau residues 244-371, C291A/P301L/ C322A/V337M) to avoid the use of reducing agents that might interfere with Clu function. Mutation of the two cysteines in TauRD avoids formation of intramolecular disulfide bonds that slows fibril formation^42^. Lipofectamine was used to render seed uptake independent of cellular machinery for internalization. The TauRD is critical for aggregation and forms the core of Tau fibrils^43, 44^.

**Fig. 1:**
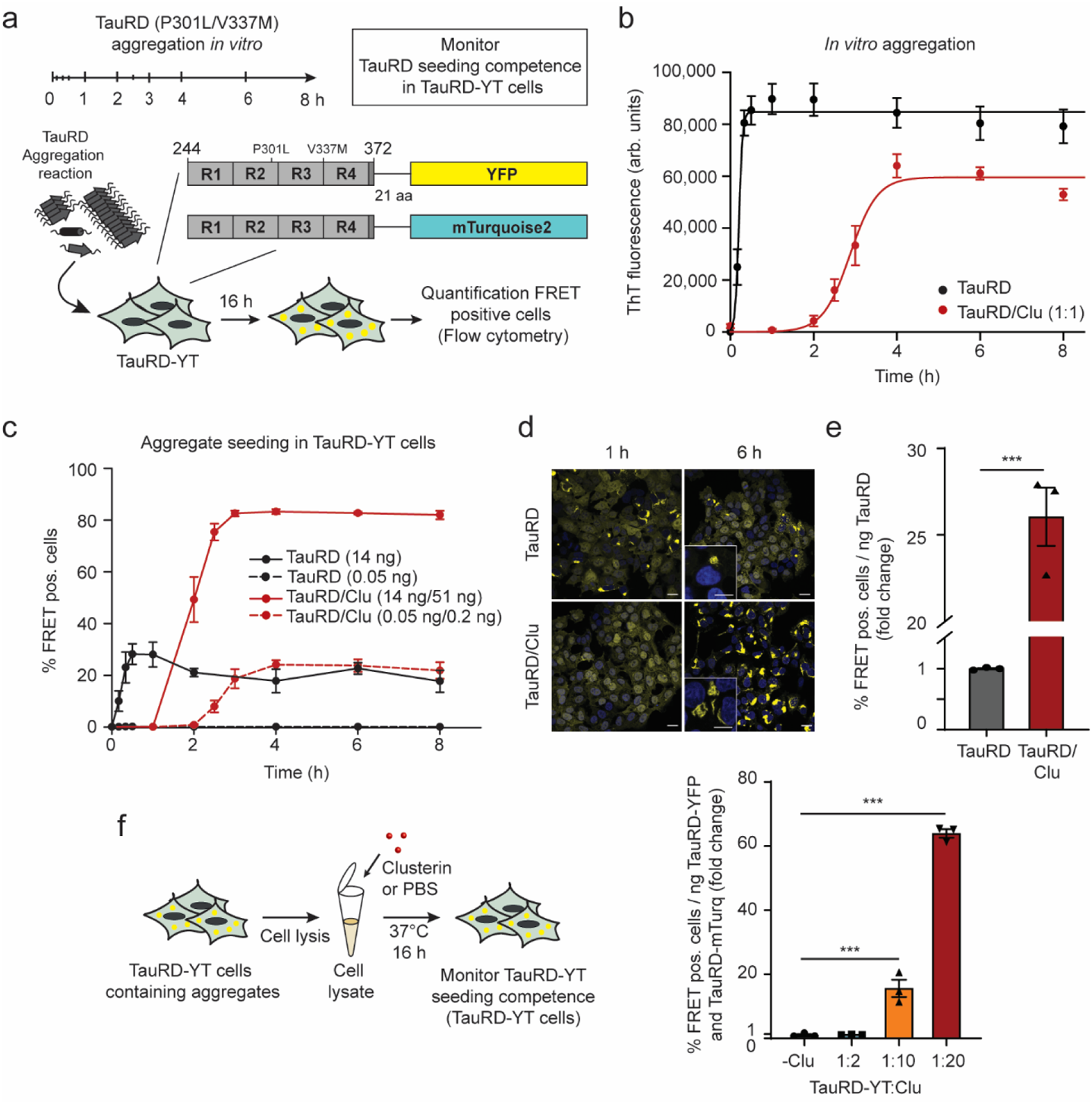
Clusterin potentiates seeding of Tau aggregates. **a**, Workflow of the seeding experiment. At the times indicated, samples were removed from TauRD aggregation reactions with or without Clu, and added with or without transfection reagent (lipofectamine) to reporter cells co-expressing TauRD fused to the FRET pair of YFP and mTurquoise2 (TauRD-YT). The fraction of cells containing FRET-positive (pos.) aggregates was quantified by flow cytometry. **b**, TauRD amyloid formation in aggregation reactions of 10 µM TauRD without (black) or with equimolar Clu (red) as monitored by ThT fluorescence. Averages ± SEM (n=10 independent experiments). arb.units, arbitrary units. **c**, Effect of Clu on formation of seeds that induce aggregation of endogenous TauRD in cells. Seed formation was analyzed as described in (a). Reporter cells were transfected with aggregation reactions containing 14 ng (solid lines) or 0.05 ng (dashed lines) TauRD and 51 ng (solid lines) or 0.2 ng (dashed lines) Clu, respectively (molar ratio Clu:TauRD 1:1). Averages ± SEM (n=3 independent experiments). **d**, Representative fluorescence microscopy images of TauRD-YT cells seeded with TauRD (14 ng TauRD) from the plateau phase of aggregation (TauRD, 1 h reaction time; TauRD/Clu, 6 h reaction time (b)). TauRD-YFP and DAPI nuclear staining signals are shown in yellow and blue, respectively. Scale bars, 20 μm for overview panels and 10 μm for insets. **e**, Fold change of seeding potency of TauRD aggregation reactions containing Clu (red) compared to control reactions without Clu (grey). Bar graphs represent the average slope ± SEM (n=3 independent experiments) from the linear regression analyses described in Supplementary Fig. 2e. *** p<0.001 (p=1.2×10^−4^) by two-tailed Student’s t-test. **f**, Effect of Clu on seeding potency of TauRD-YT aggregates contained in TauRD-YT cell lysates. Whole cell lysates of FRET-positive (pos.) TauRD-YT cells were incubated with or without Clu (molar ratios TauRD-YT:Clu 1:2, 1:10 and 1:20). Fold change of TauRD seeding potency expressed per ng of TauRD-YT in cell lysates upon treatment with increasing Clu (molar ratios TauRD-YT:Clu 1:2, 1:10 and 1:20). Bar graphs represent the average slope ± SEM (n=3 independent experiments) from the linear regression analyses shown in Supplementary Fig. 4e. ***p<0.001 (-Clu vs. TauRD-YT:Clu 1:10 p=8×10^−4^; -Clu vs. TauRD-YT:Clu 1:20 p=1.1×10^−8^) by one-way ANOVA with Bonferroni post hoc test.

TauRD rapidly formed thioflavin T (ThT)-positive fibrillar aggregates *in vitro*, induced by heparin^45^ (Fig. 1b, Supplementary Fig. 2a). Addition of Clu at an equimolar ratio to TauRD extended the lag phase and slowed fibril elongation^46, 47^ but did not prevent fibril formation (Fig. 1b, Supplementary Fig. 2a). To observe seeding, we next transferred small quantities of TauRD (0.05 ng to 14 ng with lipofectamine) after different times of *in vitro* aggregation to TauRD-YT cells, followed by analysis of endogenous aggregate formation by flow cytometry of FRET positive cells and fluorescence microscopy (Fig. 1a, c, d, Supplementary Fig. 2b). Seeding competent TauRD accumulated with kinetics similar to the formation of ThT-positive aggregates (Fig. 1b, c). The presence of Clu in the aggregation reaction delayed the appearance of seed material (0% FRET positive cells after 1 h aggregation time, Fig. 1c, d). However, once ThT-positive aggregates formed (from 2 h on, Fig. 1b), the Clu-containing aggregation reaction surprisingly presented a markedly increased seeding potency resulting in ~80% FRET-positive cells (exceeding the linear range of the assay) compared to ~30% FRET-positive cells with TauRD aggregates alone (Fig. 1c, d, Supplementary Fig. 2b). When aggregation reactions were diluted 280-fold, seeding without Clu was virtually abolished, but was still measurable in the presence of Clu, resulting in ~25% of aggregate containing cells (Fig. 1c). Again, the kinetics of seed formation correlated with the delayed formation of ThT-positive aggregates (Fig. 1b, c). The FRET-positive inclusions formed in cells with and without Clu were morphologically similar and stained with the amyloid dye X34 (Fig. 1d, Supplementary Fig. 2c). Clu alone neither formed ThT-positive species nor induced Tau aggregation when added to cells (Supplementary Fig. 2d). Titration experiments using seed material from the plateau phase of aggregation (Fig. 1b) showed that Clu increased seeding potency ~25-fold (defined as % FRET-positive cells/ng TauRD (Fig. 1e, Supplementary Fig. 2e). To test the effect of Clu on seeding in cells with unperturbed plasma membrane, we omitted the transfection reagent. Under these conditions, Clu still increased the seeding potency of TauRD aggregates ~8 fold (Supplementary Fig. 3). However, as expected, higher amounts of TauRD and TauRD/Clu aggregates were necessary to observe aggregate seeding.

The effect of Clu on TauRD aggregation and seeding is concentration dependent, since increasing the ratio of Clu relative to TauRD further delayed amyloid formation *in vitro* and increased seeding potency (Supplementary Fig. 4a, b). However, the effect on seeding potency saturated at a 1:1 molar ratio of TauRD:Clu (Supplementary Fig. 4b).

As TauRD is a highly charged protein (21 positively and 10 negatively charged amino acids), we tested whether the effect of Clu on TauRD seeding is dependent on electrostatic interactions. TauRD and TauRD/Clu aggregates (Fig. 1a) were incubated with PBS or high salt buffer (PBS/500 mM NaCl) prior to addition to TauRD-YT cells. Incubation with high salt buffer resulted in a general increase in seeding (Supplementary Fig. 4c), suggesting that high ionic strength may stabilize seeding competent TauRD species. Importantly, Clu increased seeding potency both in PBS and in the high salt buffer (Supplementary Fig. 4c), consistent with hydrophobic forces playing a role in the Clu-TauRD interaction.

In order to test the effect of Clu on preformed Tau aggregates, Clu was added to the aggregation reaction once the ThT plateau was reached (at 1 h or 24 h after initiating aggregation, Supplementary Fig. 4d). When added at an equimolar ratio to TauRD aggregates, Clu amplified seeding competence ~30-40-fold (Supplementary Fig. 4d), similar to when present during aggregation (Fig. 1c). Thus, Clu may act on preexistent fibrils or on aggregate species present in equilibrium.

The use of heparin in seed production results in a non-physiological conformation of Tau fibrils^44^. To exclude that the observed effect on TauRD seeding is dependent on heparin, we therefore investigated the effect of Clu on seeding by TauRD aggregates produced in cells. Cells were lysed under mild conditions in the presence of non-ionic detergent (0.05% Triton X-100) without sonication to preserve the structural properties of the aggregates. Lysates from TauRD-YT cells containing aggregates were incubated with increasing concentrations of Clu (Fig. 1f). A strong increase in seeding potency (up to ~60-fold) was observed upon Clu addition (Fig. 1f and Supplementary Fig. 4e). In this case higher amounts of Clu relative to TauRD (up to ~20-fold excess) were effective, presumably due to lysate proteins competing for Clu binding with the aggregates.

To exclude the possibility that our findings are limited to the isolated repeat domain of Tau, we next performed experiments with full-length Tau (FLTau 2N4R) aggregates as seeds in cells expressing either TauRD or FL-Tau FRET constructs. As expected, *in vitro* amyloid formation of FLTau was slow^42^ (t_1/2_ ~4.6 days, Fig. 2a) and was further delayed in the presence of Clu (t_1/2_ ~6.6 days) (Fig. 2a). Clu dramatically enhanced (~55-fold) the potency of FLTau aggregates to seed TauRD-YT aggregates (Fig. 2b-d and Supplementary Fig. 5a). An even greater potentiation of FLTau seeds (~100-fold) was observed with cells stably co-expressing FLTau (P301L/V337M) fusion proteins with YFP or mTuquoise2 (FLTau-YT cells), forming FLTau aggregates (Supplementary Fig. 5b-d).

**Fig. 2:**
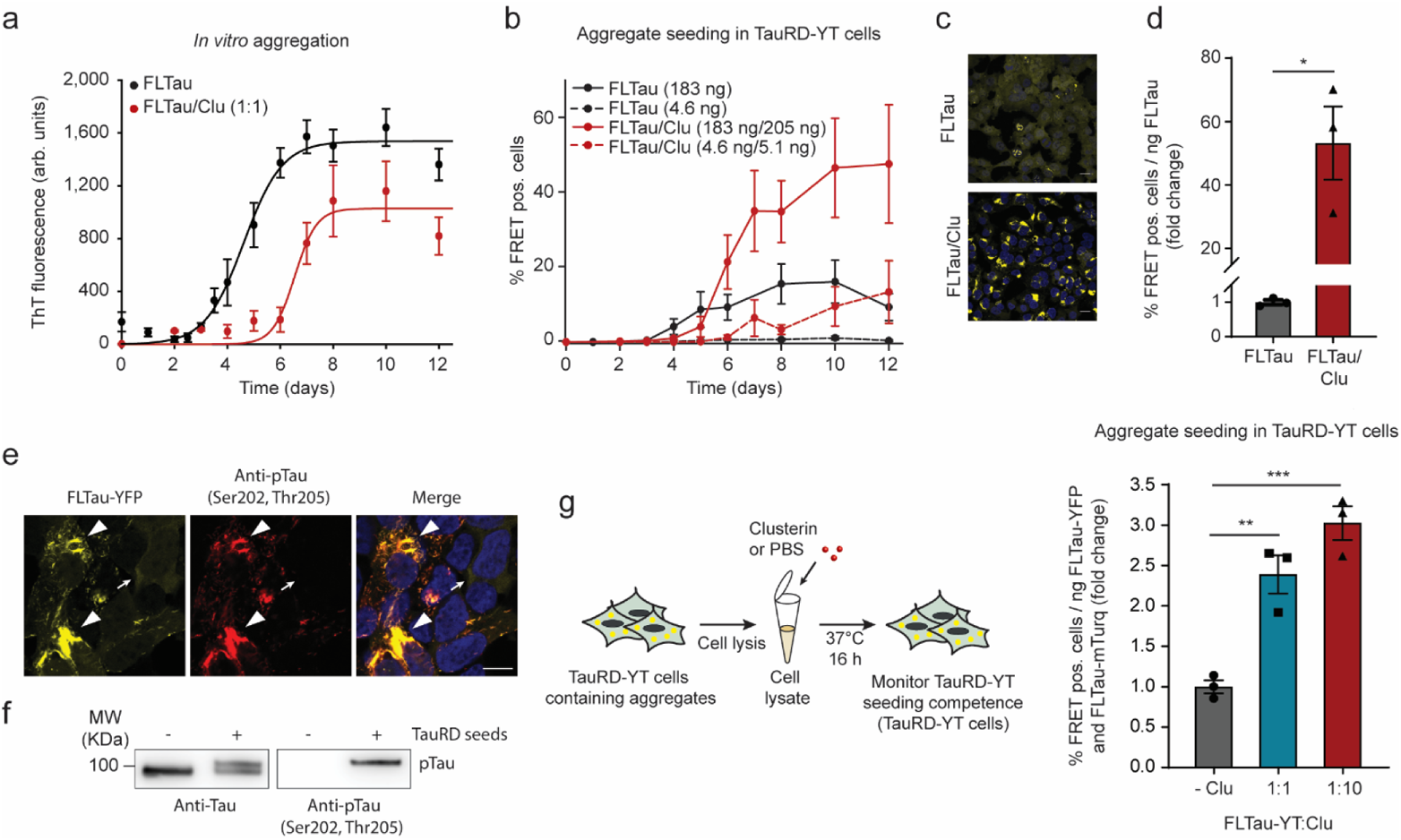
Clusterin enhances seeding potency of FLTau aggregates. **a**, Effect of Clu on kinetics of aggregation of full-length (FL) Tau (10 μM) monitored by ThT fluorescence. FLTau/Clu molar ratio was 1:1. Averages ± SEM (n=5 independent experiments). arb.units, arbitrary units. **b**, Formation of seeds in FLTau aggregation reactions without (black) or with Clu (10 μM, red) as described in (a). Reporter cells were transfected with aggregation reactions containing 183 ng (solid lines) or 4.6 ng (dashed lines) FLTau and 205 ng (solid lines) or 5.1 ng (dashed lines) Clu, respectively (molar ratio Clu:FLTau 1:1). Averages ± SEM (n=3 independent experiments). **c**, Representative fluorescence microscopy images of TauRD-YT cells seeded with FLTau aggregation reactions (183 ng FLTau) after reaching the plateau of aggregation (10 days, (a)). TauRD-YFP and DAPI nuclear staining signals are shown in yellow and blue, respectively. Scale bars, 20 μm. **d**, Fold change of seeding potency of FLTau aggregation reactions containing Clu (red) compared to control reactions without Clu (grey). Bar graphs represent the average slope ± SEM (n=3 independent experiments) from the linear regression analyses described in Supplementary Fig. 5a. * p<0.05 (p=0.0106) by two-tailed Student’s t-test. Lipofectamine was used as transfection reagent. **e, f**, Seeded aggregates of FLTau-YT contain phospho-Tau. (n=3 independent experiments) **e**, Fluorescence microscopy images of FLTau-YT cells seeded with TauRD aggregates. FLTau-YFP and immunostaining of phospho-Tau (pTau, AT8 antibody) are shown in yellow and red, respectively. The AT8 antibody recognizes Tau phosphorylation at both serine 202 and threonine 205 and is widely used to detect Tau paired helical fibrils^36, 49^. DAPI nuclear staining signal is additionally shown in blue in the merge. Arrow heads indicate aggregates. The small arrow indicates a cell without aggregates. Scale bar, 10 μm. **f**, Representative immunoblot analysis showing Tau (Tau/Repeat Domain antibody) and phospho-Tau (pTau, AT8 antibody) in FLTau-YT cell lysates from cells treated with or without TauRD seeds. Molecular weight (MW) standards are indicated. **g**, Clu enhances the seeding potency of FLTau aggregates formed in FLTau-YT cells. Whole cell lysates of FRET-positive (pos.) FLTau-YT cells were incubated without or with Clu (molar ratios FLTau-YT:Clu 1:1 and 1:10). Bar graphs represent the average slope ± SEM from the linear regression analyses described in Supplementary Fig. 5e. Data represent the mean ± SEM (n=3 independent experiments). **p<0.01 (-Clu vs. FLTau-YT:Clu 1:1 p=0.0058); ***p<0.001 (-Clu vs. FLTau-YT:Clu 1:10 p=7.9×10^−4^) by one-way ANOVA with Bonferroni post hoc test. Lipofectamine was used as transfection reagent.

As Tau aggregates in patient brain typically contain highly phosphorylated Tau^48^, we also tested the effect of Clu using cell lysates containing phosphorylated FLTau-YT aggregates as seeds^36, 49^ (Fig. 2e-g and Supplementary Fig. 5e). Phosphorylated FLTau-YT aggregates were obtained by seeding naïve FLTau-YT cells with TauRD aggregates formed *in vitro* (Fig. 1b). Phosphorylation of the resulting aggregates was confirmed by the AT8 antibody^36, 49^(Fig. 2e, f). Clu enhanced the seeding competence of these phospho-Tau aggregates up to 3-fold (Fig. 2g and Supplementary Fig. 5e). The lower effect of Clu on the seeding potency of cellular FLTau aggregates compared to TauRD aggregates (Fig. 1f and Fig. 2g) may be due to the “fuzzy coat” around the core of FLTau fibrils^50^, which may limit Clu binding, or to differential posttranslational modifications, including phosphorylation.

A physical interaction between Clu and phospho-FLTau containing aggregates was confirmed by co-immunoprecipitation using the AT8 antibody (Supplementary Fig. 5f). However, the AT8 antibody also precipitated unmodified, apparently co-aggregated FL-Tau and thus a direct interaction of Clu with phospho-Tau remains to be demonstrated.

In summary, Clu robustly enhances the potency of TauRD and FLTau aggregates to seed aggregation in cells expressing TauRD or FLTau. This effect is independent of whether Clu is present during initial aggregation or added to preformed aggregates produced *in vitro* or in cells.

### Clusterin stabilizes oligomeric Tau seeds

To biochemically characterize the seeding competent TauRD species, we fractionated the *in vitro* aggregation reaction by centrifugation. In the absence of Clu, aggregated TauRD was apparently insoluble after 1 h of aggregation (Fig. 3a, upper panel). When increasing amounts of the supernatant fraction (×10, ×20, ×30 the amount loaded in the upper panel) (Fig 3a, lower panel) were analyzed, we found ~1.5% of total TauRD to be soluble. The amount of soluble TauRD increased to ~14% of total in the presence of equimolar Clu (Fig. 3a, lower panel). Approximately 13% of Clu was recovered in the pellet fraction (Fig. 3a, upper panel), suggesting a weak association with insoluble fibrils. The seeding competence of the soluble and insoluble TauRD was compared by measuring the fraction of FRET positive cells per ng of TauRD in the seed material as determined by quantitative immunoblotting (Fig. 3a). The soluble fraction of the aggregation reaction had a much higher specific seeding capacity in the cellular assay than the resuspended pellet (Fig. 3b), indicating that soluble TauRD species are more seeding competent^51, 52^. Notably, the specific seeding activity of the Clu-containing supernatant was ~40-fold higher than that in the absence of Clu (Fig. 3b), suggesting that Clu not only increased the amount of soluble TauRD aggregates but also enhanced their intrinsic seeding activity. To test this possibility directly, we added Clu to the seeding-active, soluble fraction of a TauRD aggregation reaction generated in the absence of Clu. This resulted in a ~5-fold increase of seeding capacity (Fig. 3c). Thus, Clu not only increases the amount of soluble TauRD seeds when present during aggregation, but also enhances the intrinsic seeding activity of preexisting, soluble TauRD species.

**Fig. 3:**
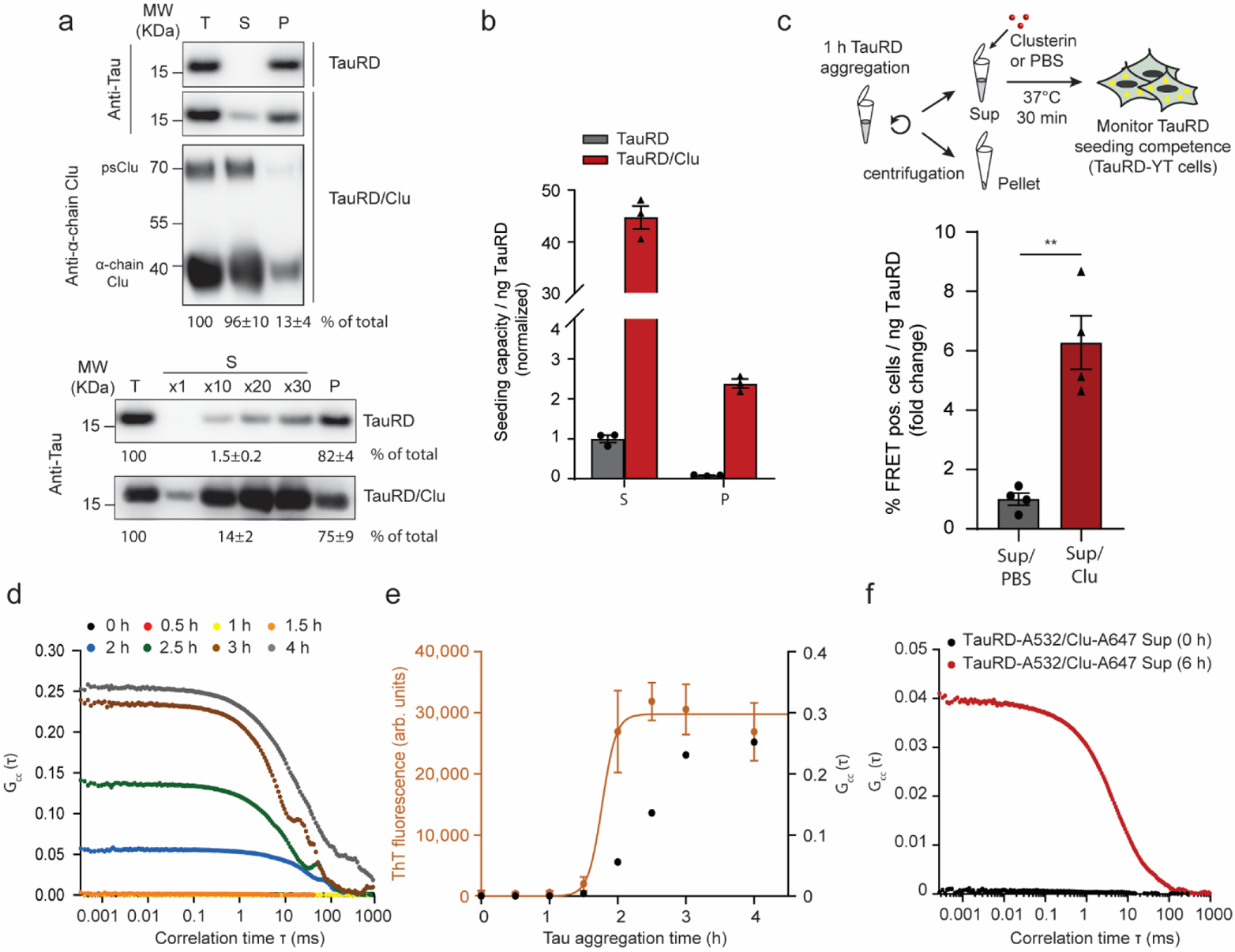
Clusterin binds to aggregates and stabilizes oligomeric Tau seeds. **a**, Fractionation by sedimentation of TauRD aggregation reactions without or with Clu (TauRD/Clu). Representative immunoblots of total (T), supernatant (S) and pellet (P) fractions. The small amount of soluble TauRD was visualized and quantified by analyzing increasing amounts of the supernatant fraction (lower panel, ×10, ×20, ×30 times the amount loaded in upper panel, x1). psClu: pre-secretory Clu. TauRD (n=6 independent experiments) and Clu (n=3 independent experiments) were quantified by densitometry with the amounts in total fractions set to 100%. Averages ± SEM. Molecular weight (MW) standards are indicated. **b**, Seeding potency of soluble (S) and pellet (P) fractions of TauRD aggregation reactions formed without (grey) or with Clu (red). Lipofectamine was used as transfection reagent. Seeding by soluble TauRD without Clu was set to 1. Averages ± SEM (n=3 independent experiments). **c**, Effect of Clu on the seeding potency of the soluble fraction of aggregation reactions without Clu. The soluble fraction was incubated for 30 min either with Clu or with PBS prior to addition to cells. Lipofectamine was used as transfection reagent. Titration of seeding potency was performed as described in Supplementary Fig. 2e. Bar graphs represent the average slope ± SEM from the linear regression analyses (n=4 independent experiments). ** p<0.01 (p=0.0013) by two-tailed Student’s t-test. **d, e**, Dual-color fluorescence cross-correlation spectroscopy (dcFCCS) analysis of the interaction of TauRD with Clu during aggregation. Aggregation reactions contained 1 μM Alexa Fluor 532 labeled TauRD (TauRD-A532) and 9 μM unlabeled TauRD in the presence of Alexa Fluor 647 labeled Clu (Clu-A647; 1.25 μM). Representative experiments are shown (n=4 independent experiments). **e**, Kinetic development of FCCS signal in (d) (right y-axis; black) relative to formation of ThT positive aggregates (left y-axis; orange, Supplementary Fig. 7a, data represent the mean ± SEM (n=3 independent experiments)). arb.units, arbitrary units. **f**, Clu interacts with soluble oligomeric TauRD species. dcFCCS analysis of TauRD-A532 and Clu-A647 interaction in the soluble fraction of the aggregation reaction immediately upon initiation of aggregation (0 h, black) and after reaching the plateau of ThT positive aggregate formation (6 h, red, Supplementary Fig. 7a). Representative data are shown (n=3 independent experiments).

Addition of Clu 6 h after initiating TauRD aggregation still produced a small, but detectable amount of soluble TauRD (Supplementary Fig. 6a), in line with enhanced seeding when Clu is added to preformed aggregates (Fig. 1f, Fig. 2g and Supplementary Fig. 4d). Surprisingly, a small amount of soluble TauRD was also generated when Clu was added to the resuspended pellet fraction (Supplementary Fig. 6b). In both cases, ~ 4-5% of total Clu associated with the insoluble TauRD aggregates (Supplementary Fig. 6a-c). Thus, Clu binding to preexistent, insoluble TauRD aggregates generates seeding competent, soluble TauRD.

To determine when during the aggregation reaction Clu interacts with TauRD, we utilized dual-color fluorescence cross-correlation spectroscopy (dcFCCS). To this end, we generated the cysteine mutant I260C in otherwise cysteine-free TauRD and labeled the protein with Alexa Fluor 532 (TauRD-A532). Residue 260 is situated outside the fibril core^44, 53^. Both Clu chains were labeled N-terminally with Alexa Fluor 647 (Clu-A647). Labeled TauRD at 1 µM was added to an aggregation reaction of unlabeled TauRD (9 µM). Labeling did not substantially affect aggregation kinetics or seeding properties of the resulting aggregates (Supplementary Fig. 7a, b, labeled TauRD; Fig. 1b, c, unlabeled TauRD) and labeled Clu (1.25 µM) enhanced seeding to a similar extent as observed with the unlabeled proteins (Supplementary Fig. 7b, labeled proteins; Supplementary Fig. 4b, unlabeled proteins; TauRD:Clu 8:1). A clear fluorescence cross-correlation signal between TauRD-A532 and Clu-A647 was only detectable at 1.5-2 h after initiation of aggregation (Fig. 3d), when ThT-positive and seeding competent TauRD species had formed (Fig. 3e). Apparently, Clu does not bind to monomeric TauRD, but interacts with TauRD aggregates. Based on their diffusion time (~16.5 ± 1.3 ms; Supplementary Fig. 7c), the TauRD/Clu complexes formed are on average ~5000 kDa in size, equivalent to Clu bound to TauRD fibrils comprising ~300 TauRD units.

To characterize the interaction of Clu with the soluble, highly seeding competent TauRD, we next measured the FCCS signal between Clu-A647 and TauRD-A532 in the soluble fraction of aggregation reactions. Clu did not interact with monomeric TauRD (0 h; Fig. 3f), but a strong interaction with soluble TauRD was detected after aggregation had reached the plateau phase (6 h; Fig 3f, Supplementary Fig. 7a). The average size of the soluble TauRD/Clu complexes was ~320 kDa (diffusion time ~4.95 ± 0.19 ms; Supplementary Fig. 7c), consistent with one or more Clu stabilizing small TauRD oligomers. The interaction between Clu and TauRD is dynamic, as the FCCS signal was rapidly reduced upon addition of excess unlabeled Clu (Supplementary Fig. 7d, e). Together these data show that Clu binds and stabilizes soluble Tau oligomers, either when present during ongoing aggregation or when added to preexistent aggregates. These soluble species are highly competent for cellular uptake and seeding of endogenous aggregation.

### Seeding competent Tau/Clu species enter cells by endocytosis

To test whether cells incorporate TauRD/Clu seeds by endocytosis, as described for Tau alone^54–58^, we incubated TauRD-YT cells during seeding with Bafilomycin, an inhibitor of the lysosomal H^+^ ATPase^59^, or leucyl-leucyl-O-methylester (LLOME), an agent that accumulates in acidic membrane compartments inducing their rupture^60^. A reduced amount of Tau/Clu aggregates compared to TauRD aggregates was applied to the TauRD-YT cells in order to obtain comparable seeding efficiencies. Both compounds increased seeding by TauRD and TauRD/Clu aggregates (in the absence of transfection reagent) to a similar extent and in a concentration dependent manner (Fig. 4a, b). Thus, seed internalization occurs via endocytosis, with a fraction of seed material presumably undergoing lysosomal degradation in the absence of Bafilomycin or LLOME.

**Fig. 4:**
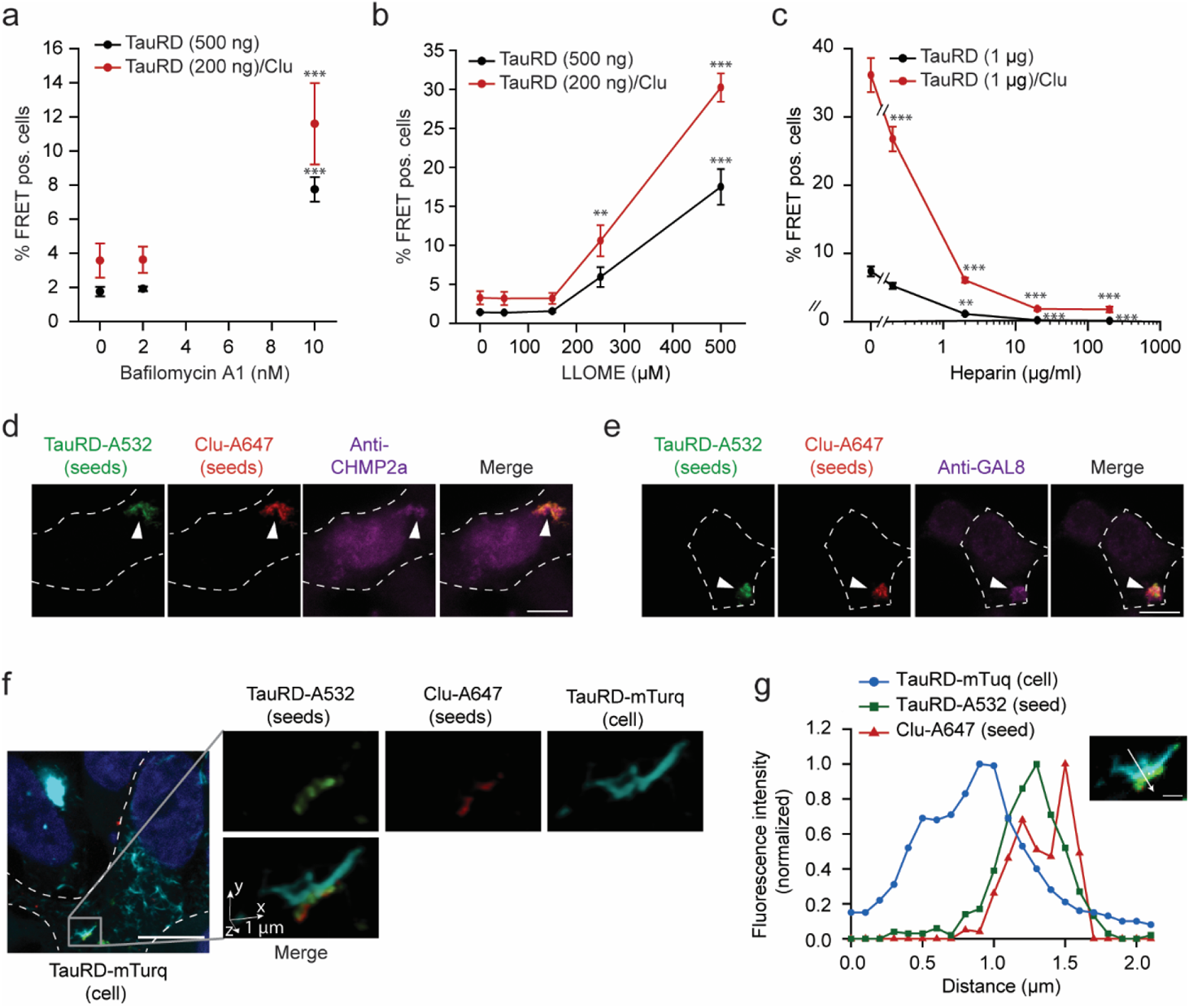
Uptake of Clusterin-associated, seeding competent Tau aggregates by endocytosis. **a, b**, Effect of inhibition of lysosomal H^+^ ATPase by Bafilomycin A1 **(a)** and permeabilization of acidic membrane compartments with LLOME **(b)** on seeding potency of TauRD aggregates formed with or without Clu. TauRD-YT cells were treated with increasing concentrations of the inhibitors in combination with 500 ng TauRD seeds or 200 ng TauRD/Clu seeds without transfection reagent. FRET positive (pos.) cells were analyzed after 48 h. Data represent the mean ± SEM (n=5 independent experiments). Significance is represented relative to control cells treated with vehicle without Bafilomycin A1 or LLOME. ** p<0.01, *** p<0.001 by two-way ANOVA with Sidak post hoc test (TauRD vehicle vs. TauRD 10 nM Bafilomycin A1 p=2.9×10^−5^; TauRD/Clu vehicle vs. TauRD/Clu 10 nM Bafilomycin A1 p=3.9×10^−4^; TauRD vehicle vs. TauRD 500 µM LLOME p=4×10^−10^; TauRD/Clu vehicle vs. TauRD/Clu 250 µM LLOME p=0.002; TauRD/Clu vehicle vs. TauRD/Clu 500 µM LLOME p<1×10^−15^). **c**, Heparan sulfate proteoglycan (HSPG) mediated internalization of TauRD and TauRD/Clu aggregates. TauRD-YT cells were treated with increasing concentrations of heparin in combination with 1 µg TauRD seeds or TauRD/Clu seeds without transfection reagent. FRET positive (pos.) cells were analyzed after 48 h. Data represent the mean ± SEM (n=4 independent experiments). Significance is represented relative to control cells treated with vehicle without heparin. ** p<0.01, *** p<0.001 by two-way ANOVA with Sidak post hoc test (TauRD vehicle vs. TauRD 2 µg/ml heparin p=0.0017; TauRD vehicle vs. TauRD 20 µg/ml heparin p=3×10^−4^; TauRD vehicle vs. TauRD 200 µg/ml heparin p=2.5×10^−4^; TauRD/Clu vehicle vs. TauRD/Clu 0.2 µg/ml heparin p=4.4×10^−6^; TauRD/Clu vehicle vs. TauRD/Clu 2 µg/ml heparin, TauRD/Clu vehicle vs. TauRD/Clu 20 µg/ml heparin and TauRD/Clu vehicle vs. TauRD/Clu 200 µg/ml heparin p<1×10^−15^). **d, e**, Colocalization of TauRD/Clu seeds (TauRD-A532 in green, Clu-A647 in red) (arrow heads) with the endocytosis marker CHMP2a (magenta) **(d)** and the marker of ruptured endomembranes, galectin-8 (GAL-8; magenta) **(e)** in TauRD-T cells. A representative result of confocal imaging is shown. The cell outline is indicated by a white dashed line. Scale bar, 10 μm. (n=3 independent experiments). **f, g**, Colocalization of TauRD/Clu seeds with endogenous TauRD-mTurquoise2 (TauRD-mTurq) aggregates. **f**, A representative slice from a confocal stack is shown (scale bar, 10 µm) where cells are outlined with a white dashed line. One aggregate region, marked with a square in the slice, is represented by volume rendering (1 µm scale bars indicated by arrows). Channels are also displayed separately. TauRD-A532 seed in green, Clu-A647 in red, endogenous TauRD aggregate in turquoise (n=3 independent experiments). **g**, Quantification of relative fluorescence intensity in the aggregate shown in the inset. TauRD-mTurq (blue), TauRD-A532 (green) and Clu-A647 (red). The colocalization line profile on a midfocal plane (inset image) along the white arrow is shown. Scale bar, 1 μm.

Heparan sulfate proteoglycans (HSPGs) are involved in cell surface binding and internalization of Tau seeds^61^, as well as in clearance of aberrant extracellular proteins mediated by Clu^31^. To test the role of this internalization mechanism in Tau/Clu seed uptake, TauRD-YT cells were incubated with increasing concentrations of heparin, a HSPG blocker, and treated with Tau aggregates formed in the presence or absence of Clu. In both cases the number of FRET positive cells strongly decreased in a manner dependent on heparin concentration (Fig. 4c), suggesting that TauRD and TauRD/Clu aggregates are internalized via HSPGs. We next investigated whether Clu and TauRD seeds enter cells together. To this end, we incubated HEK293T cells stably expressing TauRD (P301L/V337M)-mTurquoise2 (TauRD-T cells) with fluorescence labeled soluble TauRD-A532/Clu-A647 seeds (Supplementary Fig. 7a, b) for 24 h. Clu-A647 and TauRD-A532 entered the cells as a complex (Fig. 4d-g and Supplementary Fig. 8) and in several cases co-localized with CHMP2a (Fig. 4d and Supplementary Fig. 8a), a subunit of the endosomal sorting complexes required for transport (ESCRT) machinery^62^. Some seed material also co-localized with the danger receptor galectine-8 (GAL-8) (Fig. 4e and Supplementary Fig. 8b), a marker of ruptured endomembranes^63^. After incubation for further 24 h, we observed incorporation of TauRD-A532 and Clu-A647 into aggregates formed by endogenous TauRD-mTurquoise2 (TauRD-mTurq) (Fig. 4f, g and Supplementary Fig. 8c-f). The level of colocalization of the three fluorophores in seeded TauRD-mTurq aggregates was quantified by plotting their relative intensity profile, extracted from lines manually drawn in midfocal planes (Fig. 4g and Supplementary Fig, 8d, f). Co-localization of Tau-A532/Clu-A647 seeds with CHMP2a, Gal-8 and endogenous aggregates was not detected frequently enough to be reliably quantified.

Thus, following uptake by endocytosis, TauRD/Clu seeds presumably induce vesicle damage^54, 58^, allowing their escape into the cytosol where they template aggregation of endogenous TauRD.

### Clusterin interferes with seeding of α-Synuclein aggregates

To test whether the effect of Clu on aggregate seeding is Tau specific, we next performed experiments with α-Synuclein (α-Syn), which undergoes prion-like aggregate propagation in Parkinson’s disease (PD)^1^. Clu also delayed the formation of ThT-positive aggregates of α-Syn (with early-onset-PD mutation A53T), even at very low molar ratios to α-Syn^14, 64^ (Fig. 5a). However, as in the case of Tau, α-Syn fibrils nevertheless assembled (Supplementary Fig. 9a, b) and similar amounts of aggregates were generated in the absence and presence of Clu, as demonstrated by filtration assay (Supplementary Fig. 9b). Seeding competent α-Syn species formed slightly before the accumulation of ThT-positive aggregates, as analyzed in HEK293T cells stably expressing GFP-α-Syn(A53T) (Fig. 5a-c). Lipofectamine was used to render seed uptake independent of cellular machinery for internalization. In the absence of Clu, seeding with α-Syn(A53T) resulted in ~60% cells with aggregates. Seeding was markedly suppressed by substoichiometric amounts of Clu at ratios of Clu:α-Syn of 1:1,000 to 1:100 (Fig. 5b, c). This effect was also confirmed with neuroblastoma SH-SY5Y cells stably expressing GFP-α-Syn(A53T) (Supplementary Fig. 9c, d), however higher amounts of seeding material (12.5 µg α-Syn for SH-SY5Y cells, 200 ng α-Syn for HEK293T) were necessary to obtain comparable seeding efficiencies. The observed decrease in α-Syn seeding in the presence of Clu does not appear to be due to reduced formation of insoluble fibrils (Supplementary Fig. 9b), but rather to changes in the effective concentration of other seeding competent species. Note that Clu increased the seeding potency of TauRD aggregates ~2-fold even when used at a low Clu:TauRD ratio of 1:100 (Supplementary Fig. 9e). Thus, while having similar effects on aggregation kinetics *in vitro*, Clu has opposite effects on the seeding activity of Tau and α-Syn aggregates in the cellular assay, enhancing the former and suppressing the latter.

**Fig. 5:**
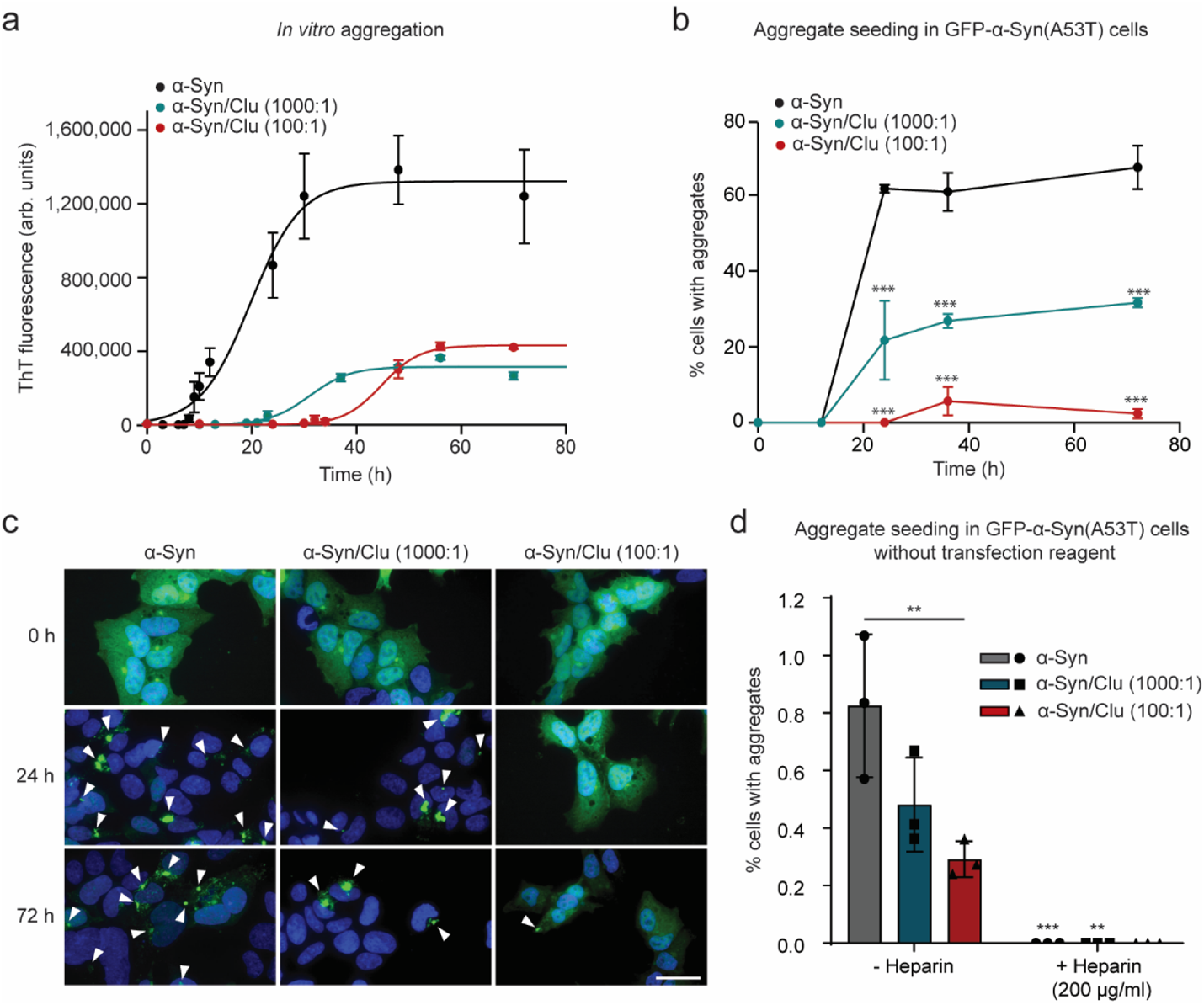
Effect of Clusterin on α-Synuclein aggregation and seeding. **a**, Aggregation of α-Syn (300 μM) without (black) or with Clu (0.3 μM and 3 μM in cyan and red, respectively) was monitored by ThT fluorescence. Molar ratios of α-Syn:Clu are indicated. arb.units, arbitrary units. Averages ± SEM (n=4 independent experiments). **b**, Effect of Clu on seeding potency of α-Syn aggregation reactions as in (a). Seeding was measured by quantifying the fraction of HEK293T cells stably expressing GFP-α-Syn(A53T) that contained aggregates after 24 h of seeding (200 ng α-Syn) with lipofectamine. Molar ratios of α-Syn:Clu are indicated. Significance is represented relative to α-Syn alone at each time point. Averages ± SEM (n=at least 100 cells examined over 3 independent experiments). ***p<0.001 by two-way ANOVA with Sidak post hoc test. (α-Syn 24 h vs. α-Syn/Clu 1000:1 24 h p=1.9×10^−8^; α-Syn 24 h vs. α-Syn/Clu 100:1 24 h p=8.4×10^−13^; α-Syn 36 h vs. α-Syn/Clu 1000:1 36 h p=4.5×10^−7^; α-Syn 36 h vs. α-Syn/Clu 100:1 36 h p=1.3×10^−11^; α-Syn 72 h vs. α-Syn/Clu 1000:1 72 h p=1.8×10^−7^; α-Syn 72 h vs. α-Syn/Clu 100:1 72 h p=2.3×10^−13^). **c**, Representative images of HEK 293T GFP-α-Syn(A53T) cells seeded with aggregation reactions (200 ng α-Syn after 0 h, 24 h and 72 h aggregation (a)) with or without Clu. GFP-α-Syn(A53T) and DAPI nuclear staining are shown in green and blue, respectively. Arrow heads indicate aggregates. Scale bar, 30 μm. **d**, Heparan sulfate proteoglycan (HSPG) mediated internalization of α-Syn and α-Syn/Clu aggregates. Seeding was measured by quantifying the fraction of GFP-α-Syn(A53T) cells that contained aggregates after 96 h of seeding (50 µg α-Syn after 72 h aggregation (a)) without lipofectamine (- Heparin). GFP-α-Syn(A53T) cells were also treated with α-Syn and α-Syn/Clu aggregates (50 µg) in combination with heparin (200 µg/ml). Molar ratios of α-Syn:Clu are indicated. Data represent the mean ± SEM (n=at least 1,000 cells examined over 3 independent experiments). ** p<0.01, *** p<0.001 by two-way ANOVA with Sidak post hoc test (-Heparin α-Syn vs. -Heparin α-Syn/Clu 100:1 p=0.003; -Heparin α-Syn vs. +Heparin α-Syn p=4.8×10^−5^; -Heparin α-Syn/Clu 1000:1 vs. +Heparin α-Syn/Clu 1000:1 p=0.007). Significance of + Heparin reactions is relative to the respective - Heparin control.

To test the effect of Clu on α-Syn seeding in cells with unperturbed plasma membrane, we omitted the transfection reagent. Under these conditions, Clu still clearly suppressed α-Syn seeding (Fig. 5d). Consistent with the notion that HSPGs are also involved in α-Syn internalization^57^, aggregate seeding by α-Syn and α-Syn/Clu was completely suppressed in the presence of heparin (200 µg/ml). Thus, HSPGs participate in the internalization of both α-Syn and TauRD seeds, independent of the presence of Clu.

### Clusterin enhances Tau seeding and toxicity in neurons

To extend our findings to neuronal cells, we used primary mouse neurons transduced with TauRD (residues 244-372, P301L/V337M) fused to YFP (TauRD-Y). We incubated neurons with either aggregates of TauRD, TauRD/Clu, or buffer control without transfection reagent and after 4 days monitored the formation of aggregates of endogenous TauRD-Y by fluorescence microscopy (Fig. 6a-c). Incubation with TauRD seeds alone induced the formation of TauRD-Y inclusions in ~12% of neurons, while TauRD/Clu induced aggregation in ~45% of neurons (Fig 6b, c). While there was no difference in neuronal viability 4 days after seeding, treatment with TauRD/Clu resulted in a ~30% decrease in viability 7 days after seeding compared to cells seeded with TauRD alone (Fig. 6d). This suggests that the higher endogenous aggregate load produced by the Clu-stabilized TauRD seeds is associated with significant neuronal toxicity.

**Fig. 6:**
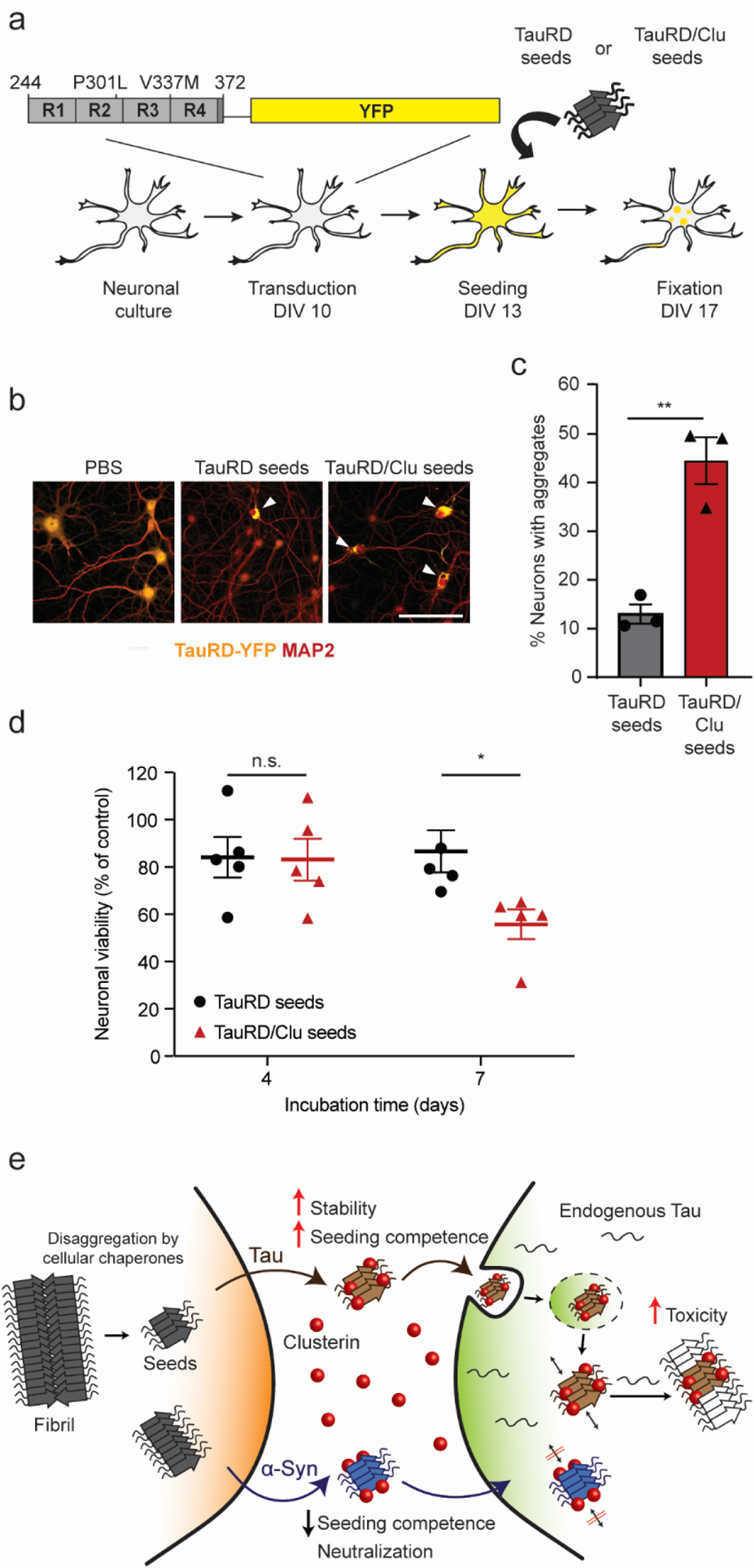
Clusterin enhances Tau seeding and toxicity in neurons. **a**, Workflow for TauRD aggregate seeding experiments with primary mouse neurons. DIV, days *in vitro*. **b**, Representative fluorescence microscopy images of primary mouse neurons expressing TauRD-YFP (yellow) incubated with PBS, TauRD seeds or TauRD/Clu seeds (70 ng TauRD). Neurons were stained with an antibody against the neuronal marker MAP2 (red). Arrow heads indicate aggregates. Scale bar, 20 μm. **c**, Comparison of seeding competence of TauRD (grey) and TauRD/Clu (red) seeds (70 ng TauRD) in primary mouse neurons. The fraction of neurons containing YFP-positive aggregates by fluorescence microscopy imaging was quantified. Data represent the mean ± SEM (n=at least 800 cells examined over 3 independent experiments). ** p<0.01 (p=0.0038) by two-tailed Student’s t-test. **d**, Viability of neurons expressing TauRD-YFP at 4 and 7 days after incubation with TauRD (black) or TauRD/Clu (red) seeds (11 ng TauRD). Data from MTT assays are normalized to the control sample incubated with PBS (100%) and represent the mean ± SEM (n=5 independent experiments). * p<0.05 (p=0.034); n.s. not significant (p=0.99) by two-way ANOVA with Sidak post hoc test. **e**, Hypothetical model for the role of Clu in amyloid seeding of Tau (brown) and α-Synuclein (α-Syn) (blue). Tau and α-Syn seeding competent species, partially produced by chaperone-mediated disaggregation, are released to the extracellular space from cells containing amyloid aggregates (grey). Clu (red) interacts with these species increasing seeding competence for Tau upon uptake by neighboring cells via the endosomal pathway. In contrast, Clu inhibits seeding for α-Syn. Tau/Clu seeds efficiently template aggregation of endogenous Tau, resulting in cytotoxicity, while α-Syn/Clu seeds are unable to template aggregation of endogenous α-Syn.

## Discussion

Extensive evidence links Clusterin with AD, and both neuroprotective and pathology-enhancing functions have been ascribed to this abundant extracellular chaperone^15–27, 31–33^. While these previous investigations focused on effects of Clu on Aβ aggregation and toxicity, our results suggest the possibility that Clu can modulate the Tau component of AD by accelerating Tau aggregate seeding. Using cellular models, we show that Clu binds and stabilizes soluble Tau species in a state highly competent in seeding aggregation of endogenous Tau upon uptake by recipient cells (Fig. 6e). The structural properties of these Tau species remain to be defined. Within the cell, clearance of aggregates by intracellular chaperones such as the Hsp70 system^65^ may generate seeding competent Tau species, which could be substrate for stabilization by Clu after release into the extracellular space (Fig. 6e). Although the exact mechanism by which Tau species are released from cells remains unclear^7^, seeding competent, high-molecular weight Tau that could be acted on by Clu, has been detected in the cerebrospinal fluid of AD patients^66^. TauRD enters target cells in complex with Clu, apparently by endocytosis (Fig. 6e). However, TauRD/Clu seeds induce vesicle rupture and escape from endosomes to the cytosol, as described for Tau alone and other amyloidogenic proteins^54, 58^(Fig. 6e), enabling them to induce aggregation of endogenous Tau. Thus, Clu fails to mediate efficient lysosomal degradation of Tau, in contrast to other Clu-bound misfolded proteins^31, 32^. Seed uptake and permeation of endolysosomal membranes may be facilitated by the relatively small size of the Tau species stabilized by Clu.

In contrast to a possible role of extracellular Clu in promoting Tau pathology, intracellular Clu, accumulating under stress conditions^9^, may be protective by interfering with *de novo* Tau aggregation. The latter activity would be consistent with a recent study reporting colocalization of Clu with intracellular Tau tangles in the brain of mice overexpressing human Tau (P301L) and exacerbated Tau pathology in CLU knock-out animals^40^. However, aggregate spreading was not explicitly assessed. It also remains to be investigated whether other apolipoproteins including ApoE, which also influences Tau pathology^67–70^, may compensate for the loss of extracellular Clu^71–74^. Whether Clu ultimately delays or promotes Tau pathology may depend on the stage of the disease and multiple other factors.

The seeding-enhancing effect of Clu appears to be Tau specific. In the case of α-Syn, another amyloidogenic protein that undergoes prion-like aggregate spreading^1^, Clu blocked both aggregation and seeding (Fig. 6e), consistent with recent observations^64, 75, 76^. Indeed, Clu is upregulated in PD and other synucleinopathies^77^. Interestingly, progressive PD in older patients is often associated with mixed brain pathologies, including Tau aggregation^78, 79^, raising the possibility of a tug of war between protective functions of Clu and collateral damage incurred. Thus, the ability of Clu to modulate transcellular Tau aggregate seeding may be broadly relevant in understanding the progressive nature of neurodegenerative pathologies.

## Methods

### Plasmids

The N1-TauRD (P301L/V337M)-EYFP construct and the mTurquoise2-N1 plasmid were gifts from Marc Diamond and Michael Davidson, respectively (Addgene #54843^80^). The N1-TauRD (P301L/V337M)-mTurquoise2 plasmid for TauRD-mTurquoise2 expression in HEK293T cells was constructed by cloning the TauRD sequence (Tau amino acids 244-372, containing the disease-related mutations P301L and V337M) from N1-TauRD (P301L/V337M)-EYFP into the mTurquoise2-N1 plasmid using the EcoRI and NheI restriction sites. The N1-FLTau (0N4R, P301L/V337M)-EYFP and N1-FLTau (0N4R, P301L/V337M)-mTurquoise2 plasmids were constructed by substitution of the TauRD (P301L/V337M) sequence from N1-TauRD (P301L/V337M)-EYFP and N1-TauRD (P301L/V337M)-mTurquoise2 plasmids by the FLTau (0N4R, P301L/V337M) sequence from the pRK5-EGFP-Tau P301L plasmid (a gift from Karen Ashe (Addgene #46908^81^) after introducing the mutation V337M, by Gibson assembly using the Gibson Assembly Master Mix (New England Biolabs).The linker between Tau and the fluorescent protein was composed of 21 amino acids in all cases (YPYGILQSTVPRARDPPVATA/M for YFP and mTurquoise2 plasmids, respectively) to avoid interference of fluorescent protein with fibril formation.

The pHUE-TauRD (P301L/V337M) plasmid was made by inserting the TauRD sequence (Tau amino acids 244-371) in pHUE^82^ by Gibson assembly using the Gibson Assembly Master Mix (New England Biolabs). Plasmid pHUE-TauRD (C291A/P301L/C322A/V337M) containing cysteine-free TauRD and pHUE-TauRD (I260C/C291A/P301L/C322A/V337M) containing TauRD-I260C for fluorescent labeling were constructed by mutagenesis of the pHUE-TauRD (P301L/V337M) plasmid using the Q5® Site-Directed Mutagenesis Kit (New England Biolabs).

The Tau/pET29b plasmid used for wild type FLTau (2N4R) expression and purification was a gift from Peter Klein (Addgene plasmid #16316^83^).

pFhSynW2 and the pVsVg packing plasmids used for lentivirus production were a gift from Dieter Edbauer. The psPAX2 packing plasmid, also used for lentivirus production, was a gift from Didier Trono (Addgene plasmid #12260). pFhSynW2 TauRD (P301L/V337M)-EYFP used for TauRD-EYFP expression in mouse primary neurons was constructed by PCR amplification of the TauRD (P301L/V337M)-EYFP sequence from the N1-TauRD (P301L/V337M)-EYFP plasmid.

The pB-T-PAF vector was a gift from James Rini. The pB-T-PAF-CluStrep plasmid, which was used for Clusterin (Clu) (Clu followed by a strep tag, WSHPQFEK) expression and purification, was constructed by cloning the Clu cDNA sequence amplified from human embryonic kidney 293T (HEK293T) cells into the pB-T-PAF vector. RNA was extracted from cell pellet using the RNeasy Mini kit (Qiagen). cDNA was then synthesized using the QuantiTect Reverse Transcription Kit (Qiagen) and the Clu cDNA amplified by PCR.The PCR product was then digested and subcloned into the pB-T-PAF vector.

Plasmid pT7-7 α-Syn A53T for the expression and purification of recombinant α-Syn was a gift from Hilal Lashuel (Addgene plasmid #105727^84^) and EGFP-α-SynA53T plasmid for α-Syn expression in HEK293T and SH-SY5Y cells was a gift from David Rubinsztein (Addgene plasmid #40823^85^)).

### Cell lines

A HEK293-EBNA suspension cell line^86^ stably expressing the recombinant protein Clusterin-Strep tag (HEK293E-CluStrep) was generated by using a piggyBac transposon-based expression system^87^ employing the pB-T-PAF-CluStrep plasmid.

The monoclonal HEK293T cell line stably expressing two different constructs, TauRD N-terminally fused to either EYFP or mTuquoise2 (TauRD-YT cell line) was generated by transfecting HEK293T cells with N1-TauRD (P301L/V337M)-EYFP and N1-TauRD (P301L/V337M)-mTurquoise2 plasmids using lipofectamine 3000 (Thermo Fisher Scientific). Cells expressing the constructs were selected by 1 mg ml^−1^ G418 antibiotic (Gibco) selection, and monoclonal cell lines were generated by isolating cells expressing both TauRD fusion proteins in a 96 well-plate by cell sorting with a BD FACS Aria III (BD Biosciences) following amplification. A monoclonal cell line allowing Tau aggregation to be monitored by flow cytometry (FRET signal detection) with high efficiency was selected. In a similar way, a monoclonal HEK293T cell line stably expressing FLTau (0N4R, P301L/V337M) fused to either EYFP or mTurquoise2 was generated (FLTau-YT cell line) using the N1-FLTau (0N4R, P301L/V337M)-EYFP and N1-FLTau (0N4R, P301L/V337M)-mTurquoise2 plasmids, as well as monoclonal cell lines expressing just one of the constructs, N1-TauRD (P301L/V337M)-EYFP construct (TauRD-Y cell line), N1-TauRD (P301L/V337M)-mTurquoise2 construct (TauRD-T cell line), N1-FLTau (0N4R, P301L/V337M)-EYFP construct (FLTau-Y cell line) and N1-FLTau (0N4R, P301L/V337M)-mTurquoise2 construct (FLTau-T cell line).

The HEK293T and SH-SY5Y cell lines stably expressing EGFP-α-Syn(A53T) ^88^ were generated by transfecting HEK293T and SH-SY5Y cells with the EGFP-α-SynA53T plasmid using lipofectamine (Thermo Fisher Scientific). Cells expressing the constructs were selected by 2000 and 1000 µg ml^−1^ G418 antibiotic (Gibco) treatment, respectively. The SH-SY5Y EGFP-α-Syn(A53T) cell line was enriched by selecting cells expressing EGFP-α-SynA53T by cell sorting with a BD FACS Aria III (BD Biosciences) following amplification.

HEK293T and SH-SY5Y cell lines were maintained at 37 °C and 5% CO_2_ in Dulbecco’s modified Eagle’s medium (DMEM, Biochrom KG) supplemented with 10% fetal bovine serum (Gibco), 100 U ml^−1^ penicillin (Gibco), 100 µI ml^−1^ streptomycin sulfate (Gibco) and 2 mM L-glutamine (Gibco). Stable cell lines were maintained in the medium described above supplemented with G418 (200 µg ml^−1^).

### Primary neuronal cultures

Primary mouse neurons were prepared from 15.5 CD-1 wild type embryos. All experiments involving mice were performed in accordance with the relevant guidelines and regulations. Pregnant females were sacrificed by cervical dislocation, the uterus was removed from the abdominal cavity and placed into a 10 cm sterile Petri dish on ice containing dissection medium, consisting of Hanks’ balanced salt solution (HBSS) supplemented with 0.01 M HEPES, 0.01 M MgSO_4_, and 1% Penicillin/Streptomycin. Each embryo was isolated, heads were quickly cut, and brains were removed from the skull and immersed in ice-cold dissection medium. Cortical hemispheres were dissected and meninges were removed under a stereo-microscope. Cortical tissue from typically six to seven embryos was transferred to a 15 ml sterile tube and digested with 0.25% trypsin containing 1 mM ethylenediaminetetraacetic acid (EDTA) and 15 μl 0.1% DNAse I for 20 minutes at 37 °C. The enzymatic digestion was stopped by removing the supernatant and washing the tissue twice with Neurobasal medium (Invitrogen) containing 5% FBS. The tissue was resuspended in 2 ml medium and triturated to achieve a single cell suspension. Cells were spun at 130 x g, the supernatant was removed, and the cell pellet was resuspended in Neurobasal medium with 2% B27 (Invitrogen), 1% L-Glutamine (Invitrogen) and 1% Penicillin/Streptomycin (Invitrogen). For immunofluorescence, neurons were cultured in 24-well plates on 13 mm coverslips coated with 1 mg ml^−1^ poly-D-Lysine (Sigma) and 1 µg ml^−1^ laminin (Thermo Fisher Scientific) (100,000 neurons per well). For viability measurements, neurons were cultured in 96-well plates coated in the same way (19,000 neurons per well). Lentiviral transduction was performed at DIV 10. Viruses were thawed and immediately added to freshly prepared neuronal culture medium. Neurons in 24-well plates received 1 μl of virus per well. Neurons in 96-well plates received 0.15 μl of virus per well. A fifth of the medium from cultured neurons was removed and the equivalent volume of virus-containing medium was added.

### Lentivirus production

HEK293T cells for lentiviral packaging (Lenti-X 293T cell line, Takara) were expanded to 70-85% confluency in DMEM Glutamax (+ 4.5g l^−1^ D-Glucose, - Pyruvate) supplemented with 10% FBS (Sigma), 1% G418 (Gibco), 1% NEAA (Thermo Fisher Scientific), and 1% HEPES (Biomol). Only low passage cells were used. For lentiviral production, a three-layered T525cm^2^ flask (Falcon) was seeded and cells were henceforth cultured in medium without G418. On the following day, cells were transfected with the pFhSynW2 TauRD (P301L/V337M)-EYFP expression plasmid and the packaging plasmids psPAX2 and pVsVg using TransIT-Lenti transfection reagent (Mirus). The transfection mix was incubated for 20 min at room temperature (RT) and in the meanwhile, cell medium was exchanged. 10 ml transfection mix were added to the flask and left overnight. The medium was exchanged on the next day. After 48-52 h, culture medium containing the viral particles was collected and centrifuged for 10 min at 1200 x g. The supernatant was filtered through 0.45 μm pore size filters using 50 ml syringes, and Lenti-X concentrator was added (Takara). After an overnight incubation at 4 °C, samples were centrifuged at 1,500 x g for 45 min at 4 °C, the supernatant was removed, and the virus pellet was resuspended in 600 µl TBS-5 buffer (50 mM Tris-HCl pH 7.8, 130 mM NaCl, 10 mM KCl, 5 mM MgCl_2_). After aliquoting, the virus was stored at −80°C.

### Chemicals and cell treatments

Bafilomycin A1 was purchased from InvivoGen. L-Leucyl-L-Leucine methyl ester (hydrochloride) (LLOME) was purchased from Cayman chemicals. Both compounds were dissolved in DMSO and small aliquots were stored at −20 °C until further use. For non-treated samples, DMSO alone was used as control. Alexa Fluor 532 C5 maleimide and Alexa Fluor 647 N-hydroxysuccinimide (NHS) ester were purchased from Thermo Fischer Scientific and freshly dissolved in DMSO before protein labeling. Heparin sodium salt from porcine intestinal mucosa was purchased from Merck (H3393).

### Sodium dodecyl sulfate-polyacrylamide gel electrophoresis (SDS-PAGE) and immunoblotting

Protein samples were boiled in SDS-PAGE sample buffer for 5 min. Protein samples were separated by electrophoresis on NuPAGE 4%–12% Bis-Tris SDS gels (Thermo Fisher Scientific) using NuPAGE MES SDS running buffer (Thermo Fisher Scientific) at 180 V. Coomassie blue staining was performed with InstantBlue (Merck). For immunoblotting, proteins were transferred at 70 V for 2 h onto a nitrocellulose membrane (GE Healthcare) using a wet electroblotting system (BIO-RAD). Membranes were blocked for at least 1 h with 0.05% TBS-Tween and 5% low fat milk. Immunodetection was performed using mouse monoclonal Tau/Repeat Domain antibody (TECAN, 2B11), anti-Tau-1 antibody (non-phosphorylated Tau, clone PC1C6, MERCK, MAB3420), Tau monoclonal antibody (TAU-5, Total Tau, Thermo Fisher Scientific, MA5-12808), phospho-Tau antibody (Ser202, Thr205, AT8, Thermo Fisher Scientific, MN1020), mouse monoclonal Clu-α antibody (Santa Cruz Biotechnology, sc-5289), recombinant anti-Clusterin antibody (EPR2911, abcam, ab92548) and anti-alpha-Synuclein antibody (LB509, abcam, ab27766). Conjugated goat-anti mouse immunoglobulin G (IgG)-horseradish peroxidase (HRP) (Merck, A4416) or anti-rabbit IgG (whole molecule)–peroxidase antibody produced in goat (MERCK, A9169) were used as secondary antibodies. Immobilon Classico Western HRP substrate or Immobilon ECL Ultra Western HRP Substrate (Merck) were used for detection. Quantification by densitometry was performed with AIDA Image Analyzer v.4.27 (Elysia Raytest) or Image J (Rasband, W.S., National Institutes of Health, USA). Full scan blots are provided in the Source Data file.

## Protein purification

Seed aggregates for addition to cells were generated with purified recombinant cysteine-free TauRD (Tau residues 244-371, C291A/P301L/C322A/V337M) to avoid the use of reducing agents that might interfere with Clu function (Supplementary Fig. 1a). Mutation of the two cysteines in TauRD avoids formation of intramolecular disulfide bonds that slows fibril formation^42^. Cysteine-free TauRD and TauRD-I260C were expressed as N-terminal His_6_-ubiquitin fusion proteins in *Escherichia coli* BL21(DE3) cells transformed with the respective pHUE-TauRD plasmids via IPTG induction. The cell pellet from a 2 l culture was resuspended in 50 ml lysis buffer (50 mM PIPES-NaOH pH 6.5, 250 mM NaCl, 10 mM imidazole, 2 mM β-mercaptoethanol (βME)) supplemented with 1 mg ml^−1^ lysozyme, Complete EDTA-free protease inhibitor cocktail (MERK) and benzonase, and incubated while gently shaking at 4 °C for 30 min. Cells were lysed by ultra-sonication, and the lysate cleared by centrifugation (1 h, 40,000 x g at 4 °C). The supernatant was loaded onto a Ni-NTA column equilibrated with lysis buffer. The column was washed with high salt buffer (50 mM PIPES-NaOH pH 6.5, 500 mM NaCl, 10 mM imidazole, 2 mM βME) and wash buffer (50 mM PIPES-NaOH pH 6.5, 250 mM NaCl, 50 mM imidazole, 2 mM βME), and His_6_-ubiquitin-TauRD was eluted with elution buffer (50 mM PIPES-NaOH pH 6.5, 50 mM NaCl, 250 mM imidazole, 2 mM β-ME). The eluted fractions were collected and the salt concentration was reduced by diluting the sample (1:5) with PIPES buffer (50mM PIPES-NaOH pH 6.5, 2 mM βME), followed by incubation with Usp2 ubiquitin protease (0.5 mg) overnight at 4 °C in order to cleave the His_6_-ubiquitin tag. The cleavage mixture was applied to a Source30S cation exchange column and the TauRD protein was eluted with a 0-500 mM NaCl gradient in PIPES buffer. The TauRD protein was further purified by size exclusion chromatography (SEC) on Superdex-75 in phosphate buffered saline (PBS). For TauRD-I260C, the buffer used for SEC contained 1 mM tris(2-carboxyethyl)phosphine (TCEP) in order to prevent the formation of disulfide bonds. Fractions containing pure protein were combined, aliquoted and flash-frozen in liquid nitrogen for storage at −80 °C.

Wild type FLTau (2N4R) was expressed in *E. coli* BL21(DE3) transformed with the Tau/pET29b plasmid via IPTG induction. The cell pellet from 6 l of culture was resuspended in 180 ml lysis2 buffer (50 mM MES-NaOH pH 6.8, 20 mM NaCl, 1 mM MgCl₂, 5 mM DTT), applied to a French press cell disruptor and subsequently boiled for 20 min. The lysate was cleared by centrifugation (1 h, 40,000 x g at 4 °C), and the supernatant was loaded onto a Source30S cation exchange column equilibrated with lysis2 buffer. The protein was eluted with a 0-500 mM NaCl gradient and further purified by SEC on Sephacryl S-200 in 20 mM MES-NaOH pH 6.8, 20 mM NaCl, 10% glycerol. Fractions containing pure protein were combined, aliquoted and flash-frozen in liquid nitrogen for storage at −80 °C.

Recombinant Clu (CluStrep) was purified as described (dx.doi.org/10.17504/protocols.io.bvvkn64w). Strep-tagged Clu was expressed and secreted by HEK293E-CluStrep cells cultured in FreeStyle 293 Expression Medium (Thermo Fisher Scientific) for 4 days. The conditioned medium was then separated from the cells by centrifugation. For chromatographic purification, the medium was first dialyzed against wash buffer (20 mM Na acetate pH 5.0). After removal of precipitate by centrifugation, the supernatant was passed over a HiTrap SP XL cation exchange column. The column was washed with 10 column volumes (CV) denaturing buffer (20 mM Na acetate pH 5.0, 6 M urea), followed by 5 CVs wash buffer. For protein elution, a 0-500 mM NaCl gradient in wash buffer was applied. Clu-containing fractions were further purified by SEC on Superdex-200 in 20 mM Na acetate pH 5.0, 100 mM NaCl, 1 mM EDTA. Fractions containing pure Clu were concentrated, and the buffer was exchanged to PBS using a Nap5 (GE Healthcare) column. Aliquots were flash-frozen in liquid nitrogen for storage at −80 °C.

Recombinant human α-Synuclein (α-Syn, A53T) was purified essentially as described^89^(dx.doi.org/10.17504/protocols.io.btynnpve). In brief, *E. coli* BL21(DE3) cells were transformed with the pT7-7 α-Syn A53T plasmid. Protein expression was induced with 1 mM IPTG for 4 h at 37 °C. Bacteria were harvested and the pellet was lysed in high salt buffer (750 mM NaCl, 50 mM Tris-HCl pH 7.6, 1 mM EDTA). The lysate was sonicated for 5 min and boiled subsequently. The boiled suspension was centrifuged, the supernatant dialyzed against 50 mM NaCl, 10 mM Tris-HCl pH 7.6 and 1 mM EDTA and purified by SEC on Superdex 200 in the same buffer. Fractions were collected and those containing α-Syn were combined. The combined fractions were applied onto an anion exchange column (MonoQ). α-Syn was purified by a gradient ranging from 50 mM to 1 M NaCl. Fractions containing α-Syn were combined and dialyzed in 150 mM KCl, 50 mM Tris-HCl pH 7.6. Aliquots were stored at −80 °C.

### Deglycosylation

Purified Clu was deglycosylated with PNGase F (glycerol free, NEB), following the manufacturer’s instructions.

### Rhodanese aggregation assay

Rhodanese (100 μM) was denatured in 6 M guanidinium-HCl, 5 mM DTT for 1 h at 25 °C and diluted 200-fold into PBS in the absence or presence of Clu (0.5 μM). Bovine serum albumin (BSA) (Thermo Fisher Scientific) (0.5 μM) was used as control. Aggregation was monitored immediately after dilution by measuring turbidity at 320 nm wavelength.

### Filter retardation assay

Different amounts of α-Syn aggregation reaction were diluted in PBS and subsequently applied onto a pre-wetted 0.2 µm pore size cellulose acetate membrane in a Hoefer slot-blot apparatus. The membrane was subsequently washed twice with 0.1 % Triton-X 100. Immunodetection was performed as described above (Sodium dodecyl sulfate-polyacrylamide gel electrophoresis (SDS-PAGE) and immunoblotting).

### Protein labeling

TauRD-I260C and Clu were labeled with Alexa532 C5 maleimide and Alexa647 NHS ester (Thermo Fisher Scientific), respectively. Before the labeling reaction, TauRD-I260C was incubated on ice in the purification buffer containing 2 mM TCEP to reduce the cysteine residue, followed by removal of TCEP by SEC using a Nap5 (GE Healthcare) column pre-equilibrated with PBS buffer. Labeling of TauRD-I260C at equimolar ratio of fluorophore was performed in the PBS buffer for 1 h at RT. For the labeling reaction of Clu, PBS buffer was exchanged with 0.1 M sodium bicarbonate buffer (pH 8.3) (N-terminal labeling buffer) using a Nap5 column and labeling was subsequently performed at a 2-fold molar excess of fluorophore for 1 h at RT. Free dyes were removed using a Nap5 column, pre-equilibrated with PBS buffer. The labeling efficiency was measured by nanodrop and was typically about 65-70%.

### Fluorescence cross-correlation spectroscopy (dcFCCS)

dcFCCS measurements were performed on a Micro Time 200 inverse time-resolved fluorescence microscope (PicoQuant) as described previously^90, 91^. Samples were diluted 100-fold in PBS (from 1 μM to 10 nM each labeled protein) immediately before each measurement. Autocorrelation was recorded for 30 min. The diffusion time was obtained by fitting the curves with the triplet diffusion equation using Symphotime 64 (PicoQuant). The confocal volume (V_eff_) was calibrated daily using Atto655 maleimide dye.

### Estimation of the average molecular weight of Tau/Clu complexes

To estimate the average size of Tau/Clu complexes, we performed FCS measurements to determine the diffusion coefficient (D). Diffusion coefficients (D) were converted into hydrodynamic radii (R_H_) via the Stokes-Einstein equation (Eq. 1), where k_B_ is the Boltzmann constant, T is the temperature (in K) and η is the solvent viscosity. Second, the correlation between R_H_ and the chain length (in amino acid residues, N) (Eq. 2) for elongated proteins was applied^92^, and this chain length was converted to molecular weight by assuming an average molar mass for amino acids of m_aa_= 113 g/mol (relative to amino acid abundance in eukaryotic proteins).

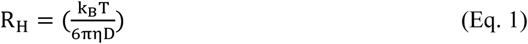

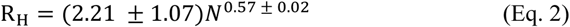

### Protein aggregation reactions and thioflavin-T (ThT) fluorescence measurements

Tau aggregation: 100 μl of 10 μM Tau (TauRD or FLTau), 2.5 μM heparin, 2 mM MgCl_2_ were incubated in the presence or absence of Clu at 37°C with constant agitation (850 rpm) in a thermomixer (Eppendorf). Aliquots were removed at the indicated time points, flash-frozen in liquid nitrogen and stored at −80 °C until measurement of ThT fluorescence and seeding activity.

α-Syn aggregation: purified α-Syn (5 mg ml^−1^, 330 μM) was centrifuged at 100,000 x g for 1 h. The supernatant was transferred into a new reaction tube and incubated in the presence or absence of Clu with constant agitation (1,000 rpm) at 37 °C in a thermomixer. Aliquots were removed at the indicated time points and stored at −80 °C until measurement of ThT fluorescence and seeding activity.

For monitoring amyloid formation by ThT fluorescence, aliquots from aggregation reactions of Tau or α-Syn were diluted 50 or 100-fold, respectively, into 20 μM ThT in PBS. Excitation and emission wavelengths were 440 nm and 480 nm, respectively. Measurements were performed with a FluoroMax-4 Spectrophotometer (HORIBA) using FluorEssence V3.9 (HORIBA). The emission signal was corrected with the reference signal of the lamp (S1/R1 wavelength, S: emission signal, R: reference signal) and by subtraction of the minimum data point. ThT kinetics were fitted using Sigma plot 14.0 (Sigmoidal dynamic fitting, sigmoid 3 parameter equation).

### Fractionation of in vitro aggregation reactions

TauRD and TauRD/Clu aggregation reactions were centrifuged at 16,100 x g and 4 °C for 1 h. The supernatant was collected. The pellet was washed with PBS, centrifuged at 16,100 x g and 4 °C for 30 min and the resulting pellet resuspended in the initial volume of PBS. The Tau and Clu content were quantified by immunoblotting.

### Cell-based seeding assays

Seeding of HEK293T cells: 100,000 cells per well of the HEK293T reporter cell line (TauRD-YT or FLTau-YT) were dispensed into a 12-well plate. For subsequent fluorescence microscopy imaging, a coverslip was placed on the well. 16-24 h later, Tau aggregates were transfected with lipofectamine 3000 (Thermo Fisher Scientific). Specifically, aggregate samples were mixed with a mixture of 50 µl Opti-MEM (Gibco) and 1.6 µl lipofectamine 3000 reagent (Thermo Fisher Scientific) and incubated for 20 min at RT. The mixtures were added to the cells with 0.5 ml of fresh medium. When lipofectamine was not used, the aggregates were mixed with 0.5 ml of fresh medium and added directly to the cells replacing the medium. After 16-20 h (when using lipofectamine) or 48 h (without lipofectamine or when using the FLTau-YT reporter cells), cells were processed for FRET signal analysis by flow cytometry or fluorescence microscopy imaging. When heparin was used to block HSPGs receptors, 1 µg of TauRD or TauRD/Clu aggregates were pre-incubated with different concentrations of heparin in 0.5 ml medium for 16 h at 4°C ^57^. After incubation, mixtures were added to the cells by exchanging the medium and 48 h later the FRET signal of seeded aggregates was analyzed by flow cytometry.

Quantification of FRET positive cells by flow cytometry: cells were harvested with TrypL Express Enzyme (Gibco), washed with PBS once and resuspended in PBS for analysis with an Attune NxT flow cytometer (Thermo Fisher Scientific). To measure the mTurquoise2 and FRET fluorescence signals, cells were excited with 405 nm laser light and fluorescence was determined using 440/50 and 530/30 filters, respectively. To measure the YFP fluorescence signal, cells were excited at 488 nm and emission was recorded using a 530/30 filter. For each sample, 50,000 single cells were analyzed. Data processing was performed using FlowJo V9 and V10.7.1 software (FlowJo LLC). After gating single cells, an additional gate was introduced to exclude YFP-only cells that show a false-positive signal in the FRET channel due to excitation at 405 nm^93^. The FRET positive gate was set by plotting the FRET fluorescence signal versus the mTurquoise2 fluorescence signal using as reference non-seeded cells (Supplementary Fig. 10).

Fluorescence microscopy imaging: cells were fixed with 4% paraformaldehyde (PFA) in PBS for 10 min, washed with PBS and permeabilized with 0.1% Triton-X100/PBS for 5 min. Nuclei were stained with NucBlue Fixed Cell ReadyProbes Reagent (Thermo Fisher Scientific) following the manufacturer’s instructions, and the coverslips were mounted with Dako fluorescence mounting medium (Agilent). Confocal imaging was performed as described below (immunofluorescence microscopy).

When lysates from cells containing aggregates were used as seeding material, cell pellets were lysed with 0.05% Triton X-100/PBS, Complete EDTA-free protease inhibitor cocktail (MERK) and benzonase for 20 min on ice, total protein was quantified by Bio-Rad Protein Assay (Bio-Rad) and the amount of Tau protein was quantified by SDS-PAGE and immunoblotting using purified Tau as standard.

Seeding of primary neurons: 70 ng of TauRD aggregates mixed with fresh medium (one tenth of medium in the well) were directly added to the neuronal cultures in 24-well plates at DIV 13. After 4 days of incubation (DIV 17), coverslips were processed as described below (immunofluorescence microscopy).

Seeding of α-Syn aggregation: HEK293T and SH-SY5Y cells expressing EGFP-α-Syn(A53T) were seeded in a 24-well plate containing a coverslip and α-Syn aggregates were transfected after 24 h using lipofectamine 2000 (Thermo Fisher Scientific). For HEK293T cells, seed material containing 200 ng of α-Syn was diluted into a mixture of 25 µl Opti-MEM (Gibco) and 1.5 µl lipofectamine. Subsequently, the suspension was added to 0.5 ml of cell culture. For SH-SY5Y cells, seed material containing 10 µg of α-Syn was diluted into a mixture of 25 µl of Opti-MEM (Gibco) and 1.5 µl of lipofectamine. Subsequently, the suspension was added to 0.5 ml of cell culture. After 24 h cells were processed for fluorescence microscopy imaging as described below and confocal imaging was performed (immunofluorescence microscopy). When heparin was used to block HSPGs receptors in seeding experiments without lipofectamine, 50 µg of α-Syn or α-Syn/Clu aggregates were pre-incubated with or without heparin (200 µg/ml) in 0.5 ml medium for 2 h at room temperature, followed by addition to the cells by medium exchange. The medium was replaced after 24 h to limit the toxicity otherwise observed with 50 µg of α-Syn. After further 48 h cells were processed for fluorescence microscopy as described below and confocal imaging was performed (immunofluorescence microscopy).

### Amyloid staining

Cells were fixed with 4% PFA/PBS for 10 min, washed with PBS and permeabilized with 0.25% Triton-X100/PBS for 30 min. Coverslips were incubated with 1 μM X-34 (Sigma) in 60% PBS, 39% ethanol, 0.02 M NaOH for 15 min, washed three times with 60% PBS, 39% ethanol, 0.02 M NaOH followed by two washes with PBS and mounted with Dako fluorescence mounting medium (Agilent).

### Immunofluorescence microscopy

Cells were fixed with 4% PFA/PBS for 10 min, washed with PBS and permeabilized with 0.1% Triton-X100/PBS for 5 min. Blocking solution (8% BSA/PBS) was added for 30 min. Coverslips were transferred to a humid chamber and incubated overnight with the primary antibody diluted in 1% BSA/PBS (anti-phospho-Tau AT8, MN1020, Thermo Fisher Scientific, 1:100 dilution; anti-CHMP2A, 10477-1-AP, Proteintech, 1:50 dilution; anti-Galectin 8, ab109519, Abcam, 1:50 dilution). Cells were then washed with 0.1% Tween-20/PBS, incubated with the corresponding secondary antibody (F(ab’)2-Goat anti-Mouse IgG Alexa Fluor 633 (A-21053); Goat-anti Rabbit IgG Alexa Fluor 405 (A-31556), Thermo Fisher Scientific) diluted in 1% BSA/PBS (1:500) for 1 h, washed with 0.1% Tween-20/PBS and stained with NucBlue Fixed Cell ReadyProbes Reagent (Thermo Fisher Scientific). Coverslips were mounted with Dako fluorescence mounting medium (Agilent). The confocal imaging was performed at the Imaging Facility of Max Planck Institute of Biochemistry, Martinsried, on a LEICA TCS SP8 AOBS confocal laser scanning microscope (Wetzlar, Germany) equipped with a LEICA HCX PL APO 63x/NA1.4 oil immersion objective. Images were analyzed with Image J (Rasband, W.S., National Institutes of Health, USA).

To detect colocalization of TauRD-A532/Clu-A647 with endogenous TauRD-mTurq aggregates, a series of z-stack images were acquired, and then deconvolved using Huygens Essentials 19.10 (Scientific Volume Imaging). Three-dimensional volume renderings were generated using Volocity V6.3 (Quorum Technologies).

Aggregate formation in primary neurons: neurons were fixed at DIV 17 with 4% PFA/PBS for 20 min; remaining free groups of PFA were blocked with 50 mM ammonium chloride in PBS for 10 min at RT. Cells were rinsed once with PBS and permeabilized with 0.25% Triton X-100/PBS for 5 min. After washing with PBS, blocking solution consisting of 2% BSA (w/v) (Roth) and 4% donkey serum (v/v) (Jackson Immunoresearch Laboratories) in PBS was added for 30 min at RT. Coverslips were transferred to a light protected humid chamber and incubated in anti-MAP2 (NB300-213, Novus Biologicals) primary antibody diluted in blocking solution (1:500) for 1 h. Cells were washed with PBS and incubated with the secondary antibody the secondary antibody Alexa Fluor 647 AffiniPure Donkey Anti-Chicken IgY (IgG) (703-605-155, Jackson Immunoresearch Laboratories) diluted 1:250 in blocking solution, with 0.5 µg ml^−1^ DAPI added to stain the nuclei. Coverslips were mounted on Menzer glass slides using Prolong Glass fluorescence mounting medium. Confocal images were obtained at a SP8 confocal microscope (Leica).

In the case of α-Syn seeding experiments, HEK293T and SH-SY5Y cells were imaged at a CorrSight microscope (Thermo Fisher) in spinning disc mode with a 63x oil immersion objective. Images were acquired with MAPS software (Thermo Fisher) and afterwards analyzed by Image J. Cells were counted manually and α-Syn accumulations with high fluorescence intensity and a diameter larger than 500 nm were considered aggregates. Fractions containing aggregates were calculated by using Origin Pro 2019b.

### Immunoprecipitation

FLTau-YT cells were lysed with 0.05% Triton X-100/PBS with Complete, Mini, EDTA-free Protease Inhibitor Cocktail (MERCK), PhosSTOP (MERCK) and benzonase for 20 min on ice. Total protein was quantified (Bio-Rad protein assay) and the amount of FLTau protein was quantified by SDS-PAGE and quantitative immunoblotting with purified FLTau as standard. Lysates were diluted in PBS (400 μl at 3 mg/ml total protein) and incubated with or without Clu (molar ratio FLTauYT:Clu 1:1) for 16 h at 37°C. Immunoprecipitation was then performed with Dynabeads Protein G Immunoprecipitation Kit (Sigma-Aldrich) following the manufacturer’s instructions. Briefly, 25 μl of magnetic beads were incubated with 5 μg of phospho-Tau antibody (AT8 antibody, MN1020, Thermo Fisher Scientific) in antibody binding and washing buffer with rotation for 30 min at room temperature. The beads were then washed and incubated with cell lysate with rotation for 1.5 h at room temperature. After incubation, the beads were washed 3 times with washing buffer, transferred to a clean tube and elution was performed by addition of 40 μl sample buffer/100 mM DTT to the beads and incubation at 70°C for 10 min. 5 μl of 1.5 M Tris pH 8.8 was added to the elution after removal of the magnetic beads. Samples (13 μl of eluate and 2 μl of lysate as input) were subsequently analyzed by SDS-PAGE and immunoblotting as described above (Sodium dodecyl sulfate-polyacrylamide gel electrophoresis (SDS-PAGE) and immunoblotting).

### Neuronal viability measurements

At DIV 13, 11 ng of TauRD aggregates mixed with fresh medium (one tenth of medium in the well) were directly added to the neuronal cultures in 96-well plates. Viability was determined with the MTT assay using thiazolyl blue tetrazolium bromide (MTT) reagent (Sigma-Aldrich). The cell medium was exchanged for 100 μl of fresh medium. Subsequently, 20 µl of 5 mg ml^−1^ MTT in PBS was added and incubated for 2-4 h at 37 °C, 5% CO_2_. Subsequently, 100 μl solubilizer solution (10% SDS, 45% dimethylformamide in water, pH 4.5) was added, and on the following day absorbance was measured at 570 nm. Each condition was measured in 6 replicates per experiment and absorbance values averaged for each experiment.

### Electron microscopy

For negative stain analysis, continuous carbon grids (Quantfoil) were glow discharged using a plasma cleaner (PDC-3XG, Harrick) for 30 s. Grids were incubated for 5 min with Tau samples, blotted and stained with 0.5% uranyl acetate solution, dried and imaged in a Titan Halo (FEI) transmission electron microscope using SerialEM.

In the case of α-Syn, grids were incubated for 1 min with the samples, blotted and subsequently washed 2 times with water for 10 s. The blotted grids were stained with 0.5% uranyl acetate solution, dried and imaged in a Polara cryo-electron microscope (FEI) at 300 kV using SerialEM.

### Statistical analysis

Statistical analysis was performed with Sigma Plot 14.0 or GraphPrism6. Sample size (n) given in figure legends describe measurements taken from distinct, independent samples. Normality was assessed in all cases. Log-transformation was applied on Bafilomycin treatment data to conform normal distribution prior statistical analysis. Two-tailed Student’s t-test was used for simple comparisons. One-way ANOVA with Bonferroni post hoc test or Two-way ANOVA with Sidak post hoc test were used for multiple comparisons.

### Data availability

All data supporting the findings of this study are available within the manuscript. Source data are provided with this paper. Other data are available from the corresponding author upon reasonable request.

## Acknowledgments

We thank Marc Diamond for the N1-TauRD (P301L/V337M)-EYFP plasmid; Michael Gropp for the pHUE-TauRD (P301L/V337M) plasmid; Saurabh Gautam for FLTau protein; Itika Saha for sharing aggregate seeding protocols; Martin Spitaler, Markus Oster and Giovanni Cardone from the Max Planck Institute of Biochemistry (MPIB) imaging facility for support with flow cytometry, imaging and image processing; Sabine Suppmann and Judith Scholz from the MPIB Biochemistry core facility for producing recombinant Clusterin; the MPIB cryo-EM facility for EM support; David Balchin for helpful discussion; Gopal Jayaraj for commenting on the manuscript and helpful discussion. The research leading to these results has received funding from the European Commission under Grant FP7 GA ERC-2012-SyG_318987–ToPAG, the Deutsche Forschungsgemeinschaft (DFG, German Research Foundation) under Germany’s Excellence Strategy within the framework of the Munich Cluster for Systems Neurology (EXC 2145 SyNergy – ID 390857198) and by the joint efforts of The Michael J. Fox Foundation for Parkinson’s Research (MJFF) and the Aligning Science Across Parkinson’s (ASAP) initiative. MJFF administers the grant ASAP-000282 on behalf of ASAP and itself. For the purpose of open access, the authors have applied a CC-BY public copyright license to the Author Accepted Manuscript version arising from this submission.

## Author contributions

F.U.H. conceived the project. P.Y. designed, performed and analyzed most of the research. V.T. conducted experiments with α-Syn. I.R. performed experiments with mouse primary neurons. R.I. conducted and analyzed dcFCCS experiments. T.S. optimized and purified recombinant Clusterin. H.W. performed Tau negative stain electron microscopy. I.D. supervised experiments with primary neurons. M.S.H. initially co-supervised the project and commented on the manuscript. A.B. co-supervised the project and contributed to experimental design. P.Y., A.B. and F.U.H. wrote the manuscript with input from the other authors.

