## Supplemental Figures for "The extracellular chaperone Clusterin enhances Tau aggregate seeding in a cellular model"

Supplementary information includes 10 Supplementary Figures.

### Supplementary Figures

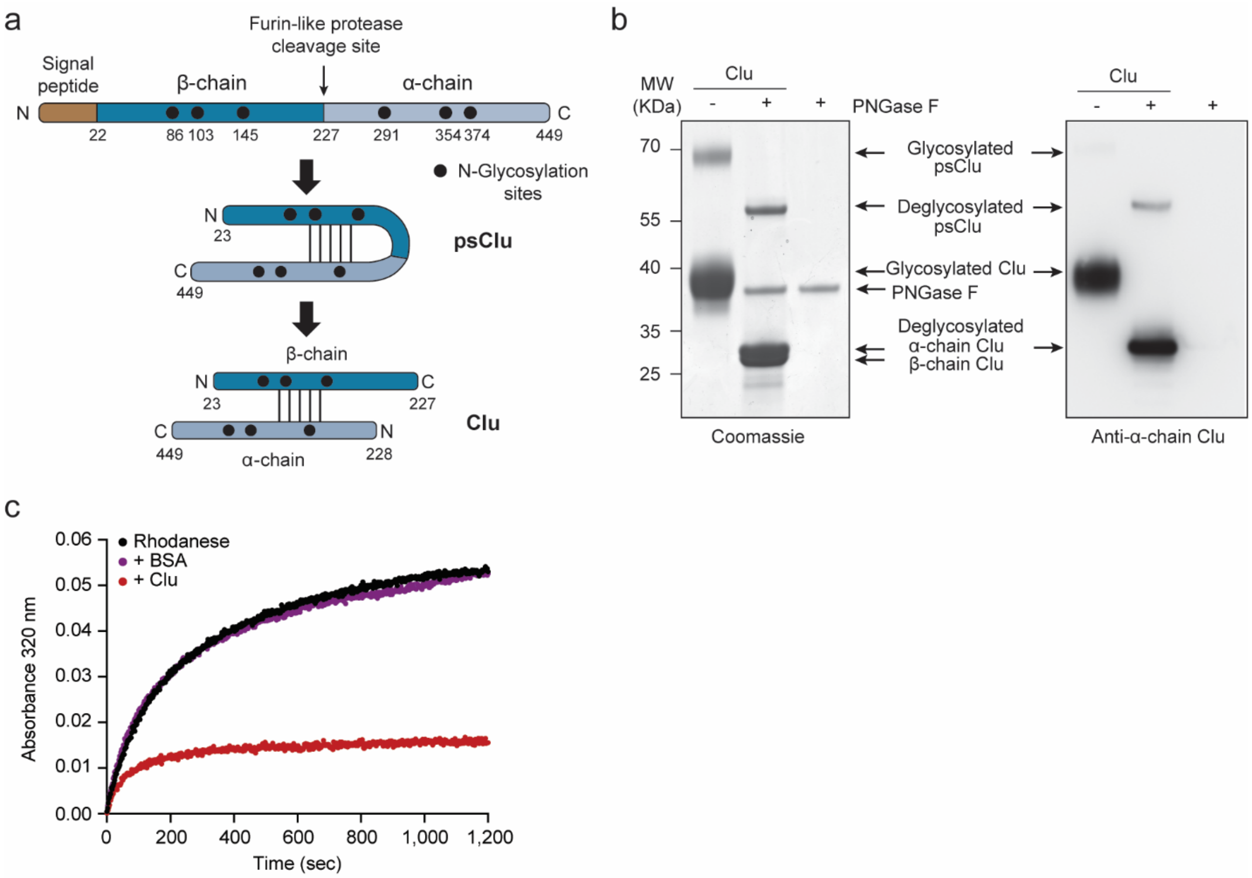


**Supplementary Fig. 1: Characterization of recombinant human Clusterin. a,** Biogenesis of Clu. The signal peptide (brown) is cleaved from the nascent chain during translocation into the ER, followed by N-glycosylation (black circles) and formation of five intramolecular disulfide bonds (black lines), resulting in pre-secretory Clu (psClu). During passage through the Golgi apparatus, psClu is processed into α- and β-chains (light and dark blue, respectively). Numbers represent amino acid position. **b,** Coomassie blue stain (left) and immunoblotting analysis (right) of SDS-PAGE gels showing recombinant human Clu before and after treatment with the glycosidase enzyme PNGase F. The SDS-PAGE samples were prepared under reducing conditions. The immunoblot shows staining with an antibody against an epitope in the α-chain of Clu. psClu: pre-secretory Clu. Molecular weight (MW) standards and protein bands are indicated. (n=3 independent experiments). **c,** Prevention of rhodanese aggregation by Clu. Bovine rhodanese denatured in 6M guanidine was 200-fold diluted (final concentration 0.5 µM) into buffer (20 mM MOPS-NaOH pH 7.2, 100 mM) containing either no protein (black), Clu (red) or bovine serum albumin (BSA, magenta) at 0.5 µM. Rhodanese aggregation at 25 °C was monitored by turbidity at 320 nm. Representative results are shown (n=3 independent experiments).

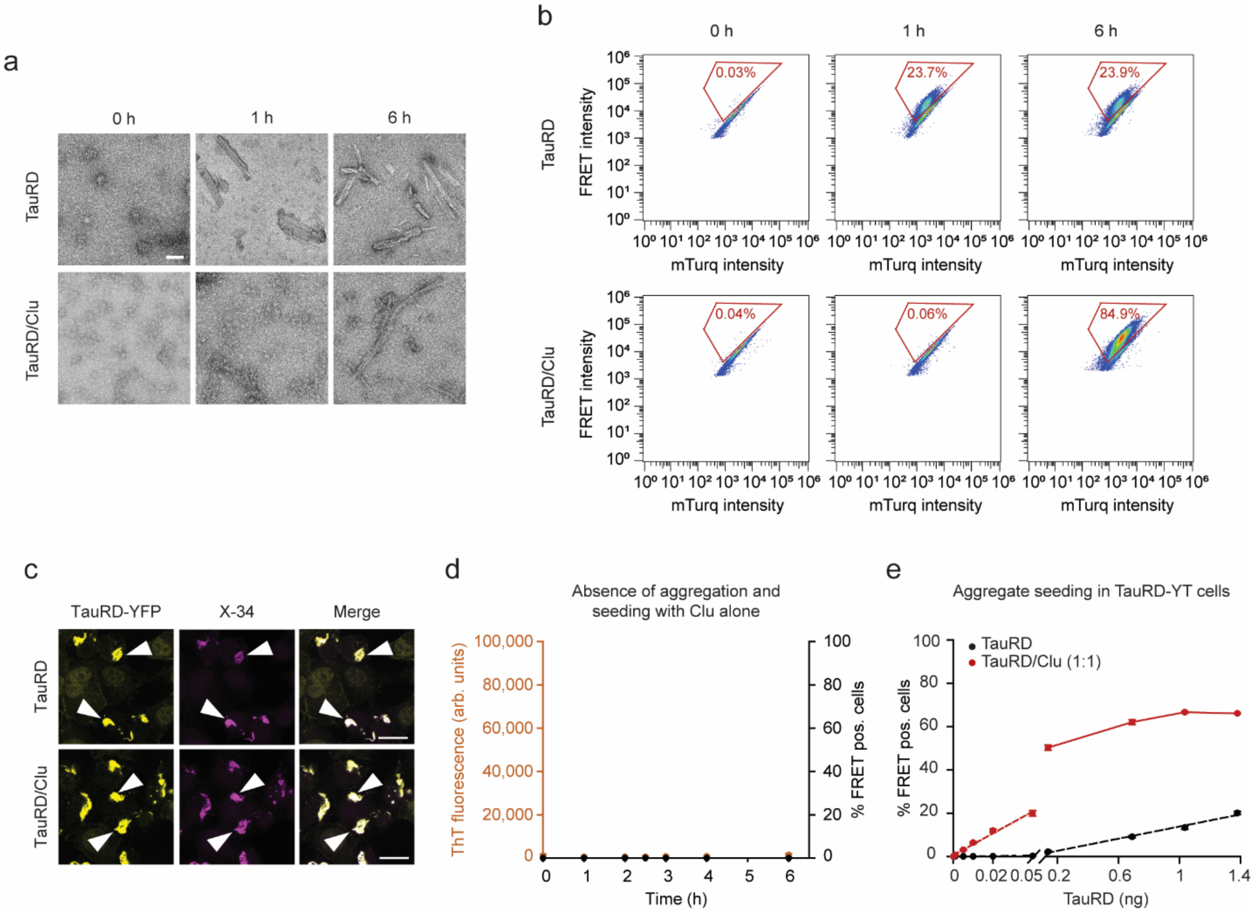


**Supplementary Fig. 2: Clusterin potentiates Tau seeding competence. a,** Negative-stain electron microscopy of TauRD aggregation reactions performed without or with equimolar Clu for the times indicated. Scale bar, 50 nm. (n=3 independent experiments). **b,** Flow cytometry analysis of TauRD-YT cells after seeding with TauRD or TauRD/Clu aggregation reactions. A double logarithmic pseudocolor dot plot representation of FRET intensity against mTurquoise2 (mTurq) intensity from individual cells is shown. Aggregation times are shown on top. The gate for "FRET-positive" cells is depicted in red and the % of FRET positive cells is indicated. **c,** Representative fluorescence microscopy images of seeded TauRD-YT cells stained with the amyloid dye X-34. Cells seeded with TauRD (top) and TauRD/Clu aggregation reactions (bottom). TauRD-YFP and X-34 staining signals are shown in yellow and magenta, respectively. Arrow heads indicate amyloid-positive TauRD-YFP aggregates. Scale bar, 20 µm. (n=3 independent experiments). **d,** Clu alone does not induce aggregate formation. Clu was incubated in the absence of TauRD and analyzed by ThT fluorescence (left y-axis; orange). After the times indicated, Clu samples were retrieved and used for seeding TauRD-YT cells (right y-axis; black). arb.units, arbitrary units. (n=3 independent experiments). **e,** Quantification of seeding potency of TauRD (black) and TauRD/Clu (red) aggregation reactions. The fraction of FRET-positive (pos.) cells after seeding is shown in dependence of the total amount of TauRD in the inoculum. Aliquots of aggregate reactions that had reached the plateau phase were used for seeding TauRD-YT cells (TauRD, 1 h reaction time; TauRD/Clu, 6 h reaction time, Fig. 1b). Seeding efficiency depends linearly on TauRD amount at low concentrations of TauRD. Dashed lines represent linear fits to the data. Data represent the mean ± SEM (n=3 independent experiments).

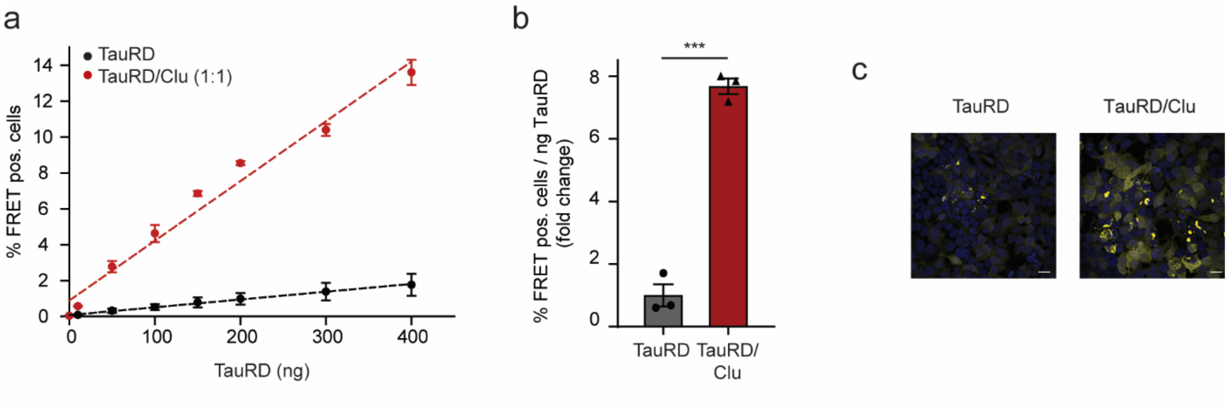


**Supplementary Fig. 3: Effect of Clusterin on TauRD seeding in the absence of transfection reagent a,** Quantification of seeding potency of TauRD (black) and TauRD/Clu (red) aggregation reactions in TauRD-YT cells in the absence of lipofectamine. Titration of seeding potency was performed as described in Supplementary Fig. 2e. Dashed lines represent linear fits to the data. Data represent the mean ± SEM (n=3 independent experiments). **b,** Fold change of seeding potency of TauRD aggregation reactions containing Clu (red) compared to control reactions without Clu (grey). Bar graphs represent the average slope ± SEM from the linear regression analyses described in (a). Data represent the mean ± SEM (n=3 independent experiments). ***$p<0.001$ ($p=1x10^{-4}$) by two-tailed Student's t-test. **c,** Fluorescence microscopy images of TauRD-YT cells seeded with 400 ng TauRD from aggregation reactions that had reached the plateau phase (TauRD, 1h reaction time; TauRD/Clu, 6 h reaction time; Fig. 1b). TauRD-YFP and DAPI nuclear staining signals are shown in yellow and blue, respectively. Scale bar, 20 µm. Experiments were performed in the absence of lipofectamine. (n=3 independent experiments).

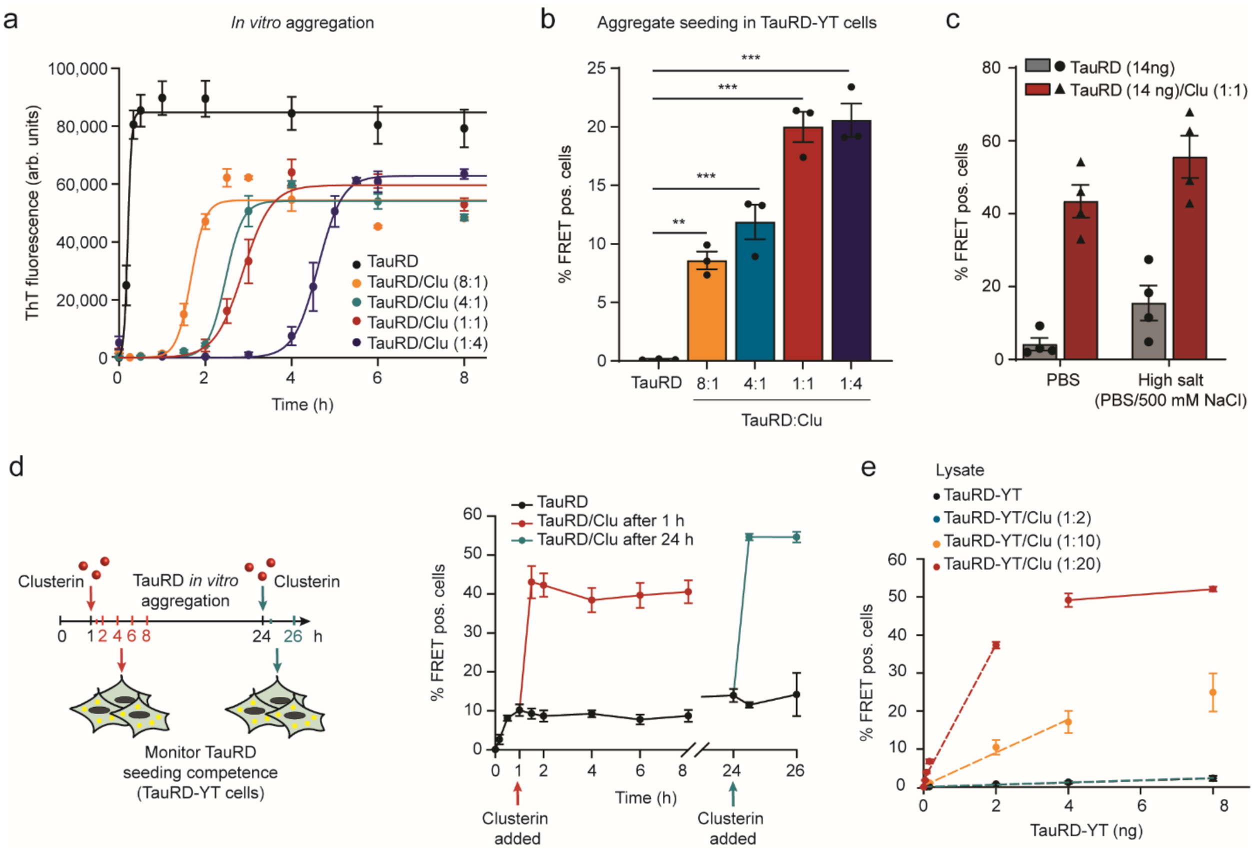


**Supplementary Fig. 4: Concentration-dependent effect of Clusterin on TauRD seeding potency, on pre-formed TauRD aggregates, and effect of ionic strength on TauRD and TauRD/Clu aggregates a,** Effect of Clu concentration on aggregation kinetics of TauRD as monitored by ThT fluorescence. Aggregation of TauRD (10 µM) without or with Clu (1.25 µM, 2.5 µM, 10 µM and 40 µM). Molar ratios of TauRD:Clu are indicated. Data represent the mean ± SEM (n=3 independent experiments). TauRD alone and TauRD:Clu equimolar ratio (n=10 independent experiments). arb.units, arbitrary units. **b,** Dependence of TauRD seeding capacity on Clu concentration in the aggregation reaction. Samples containing 0.05 ng TauRD from TauRD (10 µM) aggregation reactions with increasing concentrations of Clu (1.25 µM, 2.5 µM, 10 µM and 40 µM) were retrieved after reaching the plateau phase (1 h, 4 h, 6 h, 6 h and 8 h, respectively(a)) and used for seeding TauRD-YT cells. Molar ratios of TauRD:Clu are indicated. Lipofectamine was used as transfection reagent. Data represent the mean ± SEM (n=3 independent experiments). *** p<0.001; ** p<0.01 by one-way ANOVA with Bonferroni post hoc test. (TauRD vs. TauRD:Clu 8:1 p=0.0035; TauRD vs. TauRD:Clu 4:1 p=2.5x10$^{-4}$; TauRD vs. TauRD:Clu 1:1 p=2.2x10$^{-6}$; TauRD vs. TauRD:Clu 1:8 p=1.7x10$^{-6}$). **c,** Effect of ionic strength on TauRD and TauRD/Clu (14 ng TauRD, molar ratio TauRD:Clu 1:1) aggregate seeding. Aggregates (TauRD, 1h reaction time; TauRD/Clu, 6 h reaction time, Fig. 1b) were incubated for 1 h with PBS or high salt buffer (PBS/500 mM NaCl) prior to addition to TauRD-YT cells. Lipofectamine was used as transfection reagent. Data represent the mean ± SEM (n=4 independent experiments). **d,** Formation of seeds in TauRD (10 µM) aggregation reactions in the absence of Clu (black) or upon addition of Clu (10 µM) at 1 h (red) or 24 h (cyan) after starting aggregation. The fraction of FRET-positive (pos.) cells was monitored after seeding (14 ng

TauRD) with lipofectamine. Data represent the mean ± SEM (n=3 independent experiments). **e,** Titration experiment of the effect of Clu on seeding potency of TauRD-YT aggregates contained in TauRD-YT cell lysates. Whole cell lysates of FRET-positive (pos.) TauRD-YT cells were incubated with or without Clu (molar ratios TauRD-YT:Clu 1:2, 1:10 and 1:20, Fig. 1f). Titration of seeding potency was performed as described in Supplementary Fig. 2e. Dashed lines represent linear fits to the data. Data represent the mean ± SEM (n=3 independent experiments).

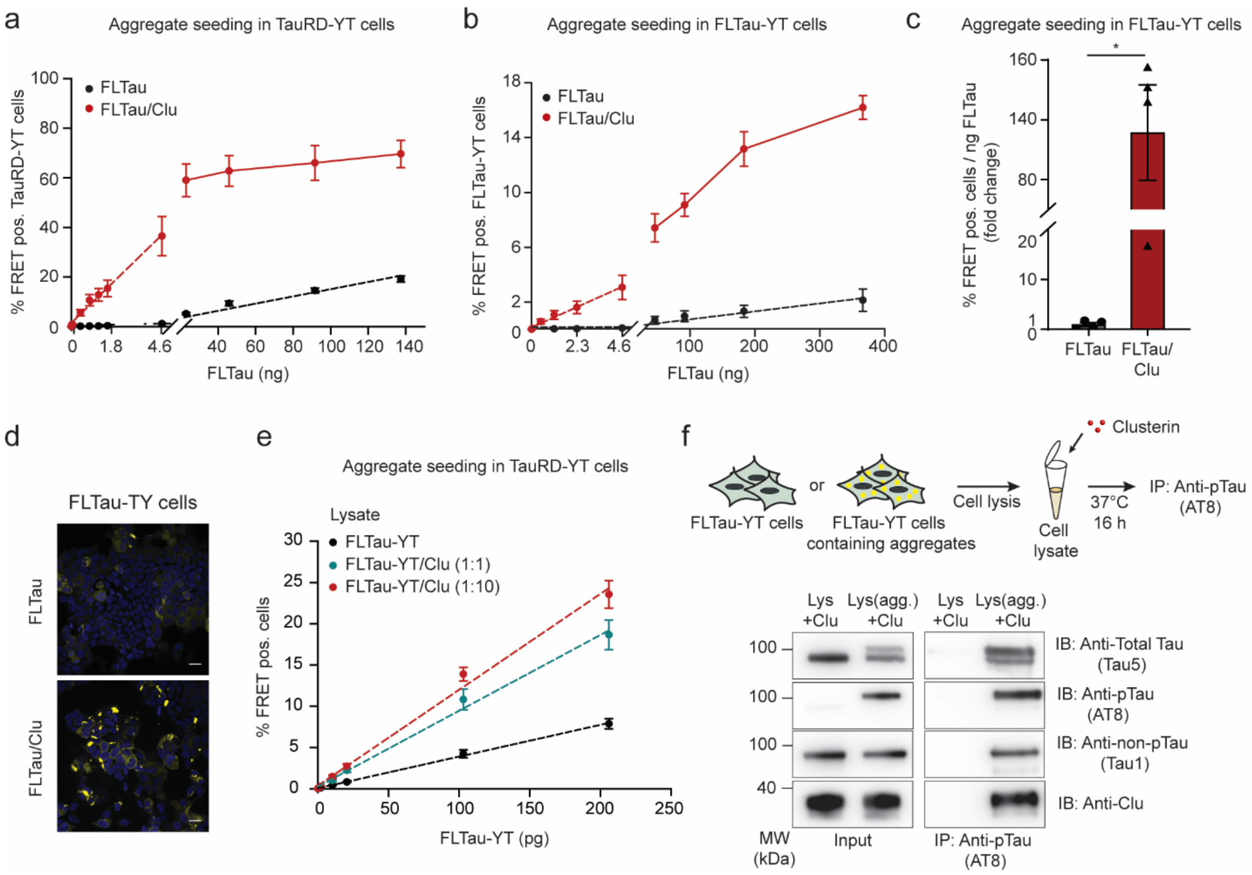


**Supplementary Fig. 5: Clusterin potentiates FLTau seeding competence and binds to aggregates containing phosphorylated FLTau. a,** Quantification of seeding potency of FLTau (black) and FLTau/Clu (red) aggregation reactions in TauRD-YT cells. Aggregate seeds were from the plateau phase of aggregation reactions (10 days, Fig. 2a). Titration of the samples was performed as described in Supplementary Fig. 2e. Dashed lines represent linear fits to the data. Data represent mean ± SEM (n=3 independent experiments). **b,** Quantification of seeding potency of FLTau (black) and FLTau/Clu (red) aggregation reactions in FLTau-YT cells. Aggregate seeds were from the plateau phase of aggregation reactions (12 days, Fig. 2a). Titration of the samples was performed as described in Supplementary Fig. 2e. Dashed lines represent linear fits to the data. Data represent mean ± SEM (n=4 independent experiments). **c,** Effect of Clu on seeding potency of FLTau aggregation reactions in FLTau-YT cells. Bar graphs represent the average slope ± SEM (n=4 independent experiments) from the linear regression analyses described in (b). *$p<0.05$ ($p=0.0134$) by two-tailed Student's t-test. **d,** Fluorescence microscopy images of FLTau-YT cells seeded with aggregation reactions (370 ng FLTau) with and without Clu. FLTau-YFP and DAPI nuclear staining signals are shown in yellow and blue, respectively. Scale bars, 20 µm. (n=3 independent experiments). **e,** Clu enhances the seeding potency of FLTau aggregates formed in FLTau-YT cells. Whole cell lysates of FRET-positive FLTau-YT cells were incubated without or with Clu (molar ratios FLTau-YT:Clu 1:1 and 1:10, Fig. 2g). Titration of the samples was performed as described in Supplementary Fig. 2e with TauRD-YT cells as recipient cells. Dashed lines represent linear fits to the data. Data represent the mean ± SEM (n=3 independent experiments). **f,** Clu binds to Tau aggregates containing phospho-FLTau-YT. Whole cell lysates of FRET-positive FLTau-YT cells (Lys(agg.)) and naïve

FLTau-YT cells (Lys) as control were incubated with Clu (molar ratio FLTau-YT:Clu 1:1) followed by p-Tau (AT8 antibody) immunoprecipitation (IP) and immunoblotting (IB) using anti-Tau5 (total Tau antibody), anti-Tau1 (non-pTau antibody), anti-AT8 (pTau antibody) and anti-Clu antibody (n=3 independent experiments). Molecular weight (MW) standards are indicated.

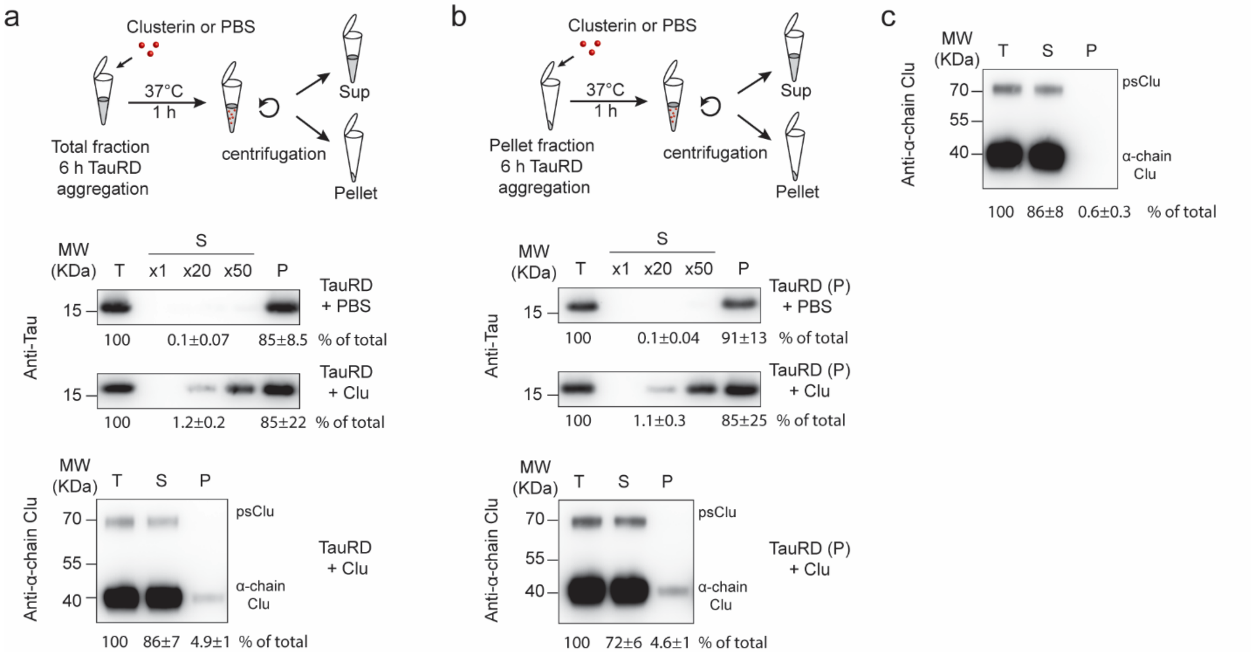


**Supplementary Fig. 6: Clusterin mediates production of soluble TauRD from pre-formed TauRD aggregates. a,** Effect of Clu on TauRD solubility when added to total aggregation reactions. TauRD aggregation reactions that had reached the plateau phase (6 h aggregation, Fig. 1b) were incubated either with PBS or Clu for 1 h at 37ºC. Reactions were fractionated by centrifugation. Total (T), supernatant (S) and pellet (P) fractions were analyzed by immunoblotting with antibodies against Tau (top panel) and α-chain Clu (bottom panel). psClu: pre-secretory Clu. TauRD and Clu were quantified by densitometry and amounts expressed as % of total. 20- (x20) and 50-fold (x50) amounts of supernatant were analyzed to quantify soluble TauRD. **b,** Effect of Clu on TauRD solubility when added to the insoluble fraction of aggregation reactions. Insoluble TauRD was isolated by centrifugation from an aggregation reaction that had reached the plateau phase (6 h aggregation, Fig. 1b). Insoluble TauRD was resuspended in PBS, followed by incubation with or without Clu for 1 h at 37ºC. Reactions were then fractionated and total (T), supernatant (S) and pellet (P) fractions analyzed as in (a). TauRD and Clu were quantified by densitometry and amounts expressed as % of total. **c,** Clu remains soluble in the absence of TauRD. Clu was incubated for 1 h at 37ºC as in (a) and (b) but in the absence of TauRD. The reaction was fractionated by centrifugation and analyzed by immunoblotting with anti-α-chain Clu antibody. Clu was quantified by densitometry and amounts expressed as % of total. Values represent mean ± SEM (n=3 independent experiments). Molecular weight (MW) standards are indicated.

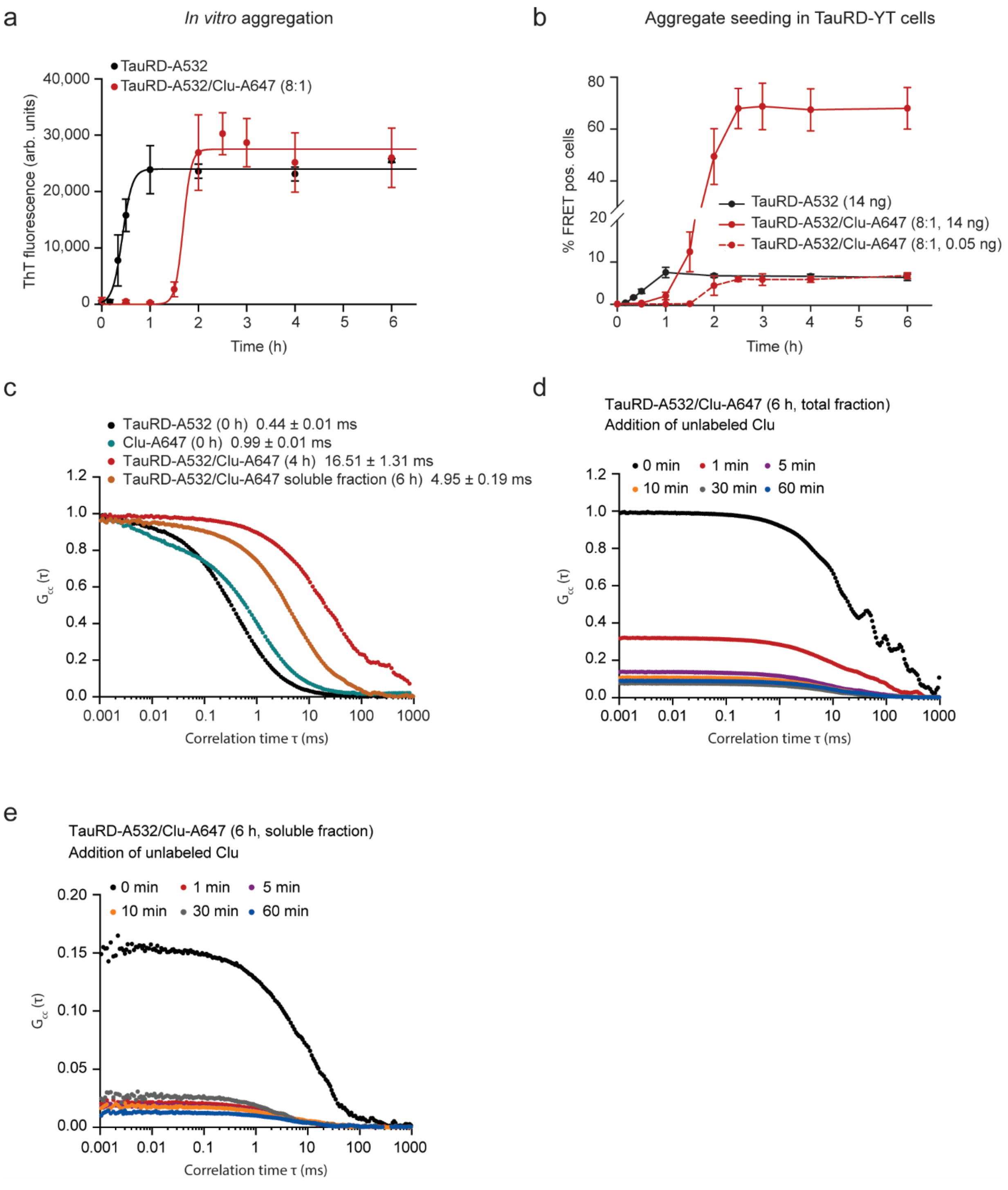


**Supplementary Fig. 7: Aggregation and seeding properties of fluorescent labeled TauRD in the presence of fluorescent labeled Clu. a,** Fluorescent labeled Clu delays aggregation of fluorescent labeled TauRD. Aggregation of TauRD-A532 (1 µM TauRD-A532 and 9 µM unlabeled TauRD) without (black) or with Clu-A647 (1.25 µM, red) was monitored by ThT fluorescence. Data represent the mean ± SEM (n=3 independent experiments). arb.units, arbitrary units. **b,** Fluorescent labeled Clu amplifies the seeding potency of fluorescent labeled

TauRD aggregates. Aggregation reactions as in (a) were used to seed TauRD-YT cells with lipofectamine as described in Fig. 1a. TauRD-YT cells were transfected with reactions containing 14 ng (solid lines) or 0.05 ng (dashed lines). Data represent the mean ± SEM (n=3 independent experiments). **c,** Diffusion times of TauRD/Clu complexes. Fluorescence correlation spectroscopy (FCS) of TauRD-A532 (black) and Clu-A647 (cyan) immediately upon initiation of aggregation (0 h). Dual-color fluorescence cross-correlation spectroscopy (dcFCCS) of the interaction of TauRD-A532 and Clu-A647 in the total aggregation reaction after 4 h of aggregation ((a), red) and dcFCCS analysis of the interaction of TauRD-A532 and Clu-A647 in the soluble fraction after 6 h of aggregation (a, orange). Representative measurements are shown. Diffusion time ± SEM (n=4 independent experiments). **d, e,** Dynamic nature of TauRD/Clu complexes in total aggregation reactions (**d**) and in the soluble fraction of aggregation reactions (**e**). Analysis of the interaction of TauRD-A532 and Clu-A647 by dcFCCS after 6 h of aggregation either without or with addition of a 10-fold excess of unlabeled Clu and incubation for 1 min to 60 min as indicated. Representative measurements are shown (n=3 independent experiments).

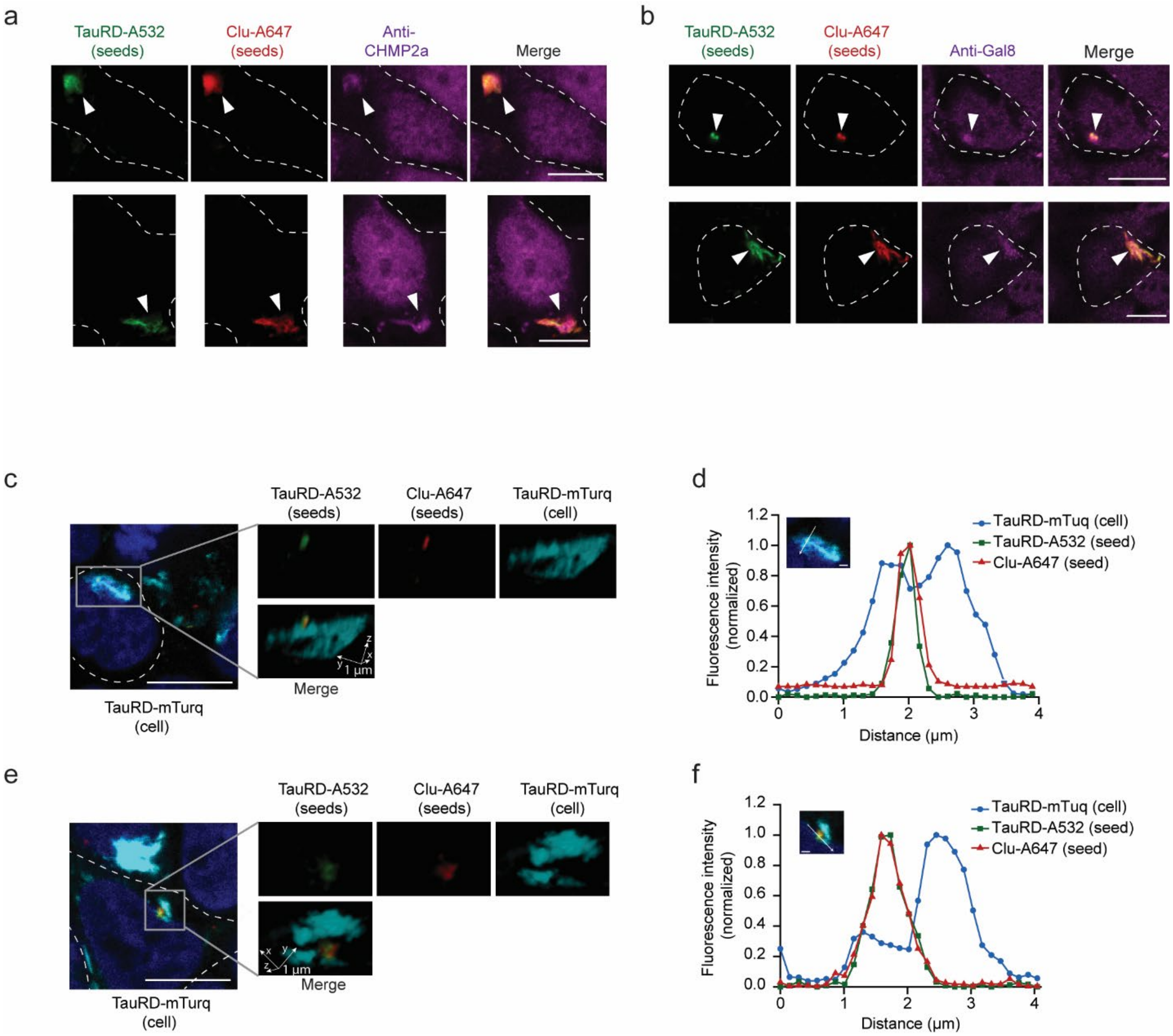


**Supplementary Fig. 8: Uptake and seeding of Clusterin-associated Tau aggregates. a, b,** Colocalization of TauRD/Clu seeds (TauRD-A532 in green, Clu-A647 in red) (arrow heads) with the endocytosis marker CHMP2a (magenta) **(a)** and the marker of ruptured endomembranes, galectin-8 (GAL-8; magenta) **(b)** in TauRD-T cells. Representative results of confocal imaging are shown. The cell outline is indicated by a white dashed line. Scale bar, 10 µm. (n=3 independent experiments). **c-f,** Colocalization of TauRD/Clu seeds with endogenous TauRD-mTurquoise2 (TauRD-mTurq) aggregates. **c, e,** Representative slices from a confocal stack are shown (scale bar, 10 µm) where cells are outlined with a white dashed line. One aggregate region, marked with a square in the slice, is represented by volume rendering (1 µm scale bars indicated by arrows). Channels are also displayed separately. TauRD-A532 seed in green, Clu-A647 in red, endogenous TauRD aggregate in turquoise. (n=3 independent experiments). **d, f,** Quantification of relative fluorescence intensity in the aggregate shown in the inset in (c) and (e), respectively. TauRD-mTurq (blue), TauRD-A532 (green) and Clu-A647 (red). The colocalization line profile on a midfocal plane (inset image) along the white arrow is shown. Scale bar, 1 µm.

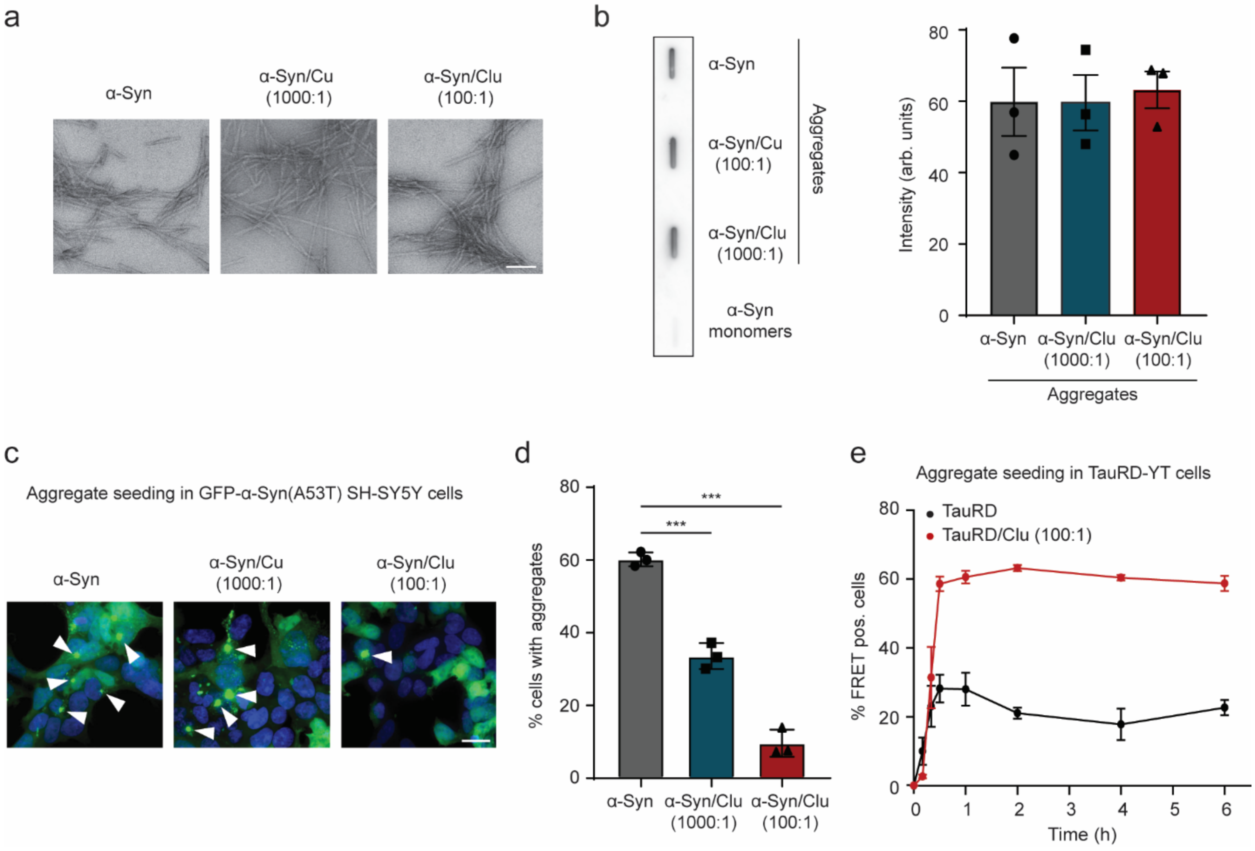


**Supplementary Fig. 9: Effect of Clusterin on α-Synuclein aggregation and seeding in SH-SY5Y cells. a,** Negative-stain electron microscopy of α-Syn aggregation reactions without or with Clu (molar ratio α-Syn:Clu 1000:1 and 100:1). Samples were analyzed after reaching the plateau phase of aggregation (72 h, (Fig. 5a)). Scale bar, 200 nm. (n=3 independent experiments). **b,** Filter retardation assay of α-Syn monomers and aggregation reactions without and with Clu (molar ratio α-Syn:Clu 1000:1 and 100:1) after reaching the plateau phase of aggregation (72 h, (Fig. 5a)). Bar graphs represent quantification by densitometry of the aggregation reactions. Average ± SEM (n=3 independent experiments). arb. units, arbitrary units. **c,** Representative images of the effect of Clu on seeding potency of α-Syn aggregation reactions as in Fig. 5a using SH-SY5Y as reporter cell line. SH-SY5Y GFP-α-Syn(A53T) cells were seeded (with lipofectamine) with aggregation reactions with or without Clu (12.5 µg α-Syn after 72 h aggregation, Fig. 5a) and cells with aggregates were analyzed after 24 h. Molar ratios of α-Syn:Clu are indicated. GFP-α-Syn(A53T) and DAPI nuclear staining are shown in green and blue, respectively. Arrow heads indicate aggregates. Scale bar, 20 µm. **d,** Bar graphs represent the quantification of the % of cells presenting aggregates after seeding as described in (c). Averages ± SEM (n=3 independent experiments). ***$p<0.001$ by one-way ANOVA with Bonferroni post hoc test. (α-Syn vs. α-Syn/Clu 1000:1 $p=1.5\times10^{-4}$; α-Syn vs. α-Syn/Clu 100:1 $p=3.5\times10^{-6}$) **e,** Substoichiometric amounts of Clu enhance seeding competence of TauRD aggregates. Seed formation was analyzed in aggregation reactions containing 10 µM TauRD without (black) or with 0.1 μM Clu (red) as judged by the fraction of FRET-positive (pos.) TauRD-YT cells. Reporter cells were transfected with aggregation reactions containing 14 ng TauRD with lipofectamine. The molar ratio of TauRD:Clu was 100:1. Averages ± SEM (n=3 independent experiments).

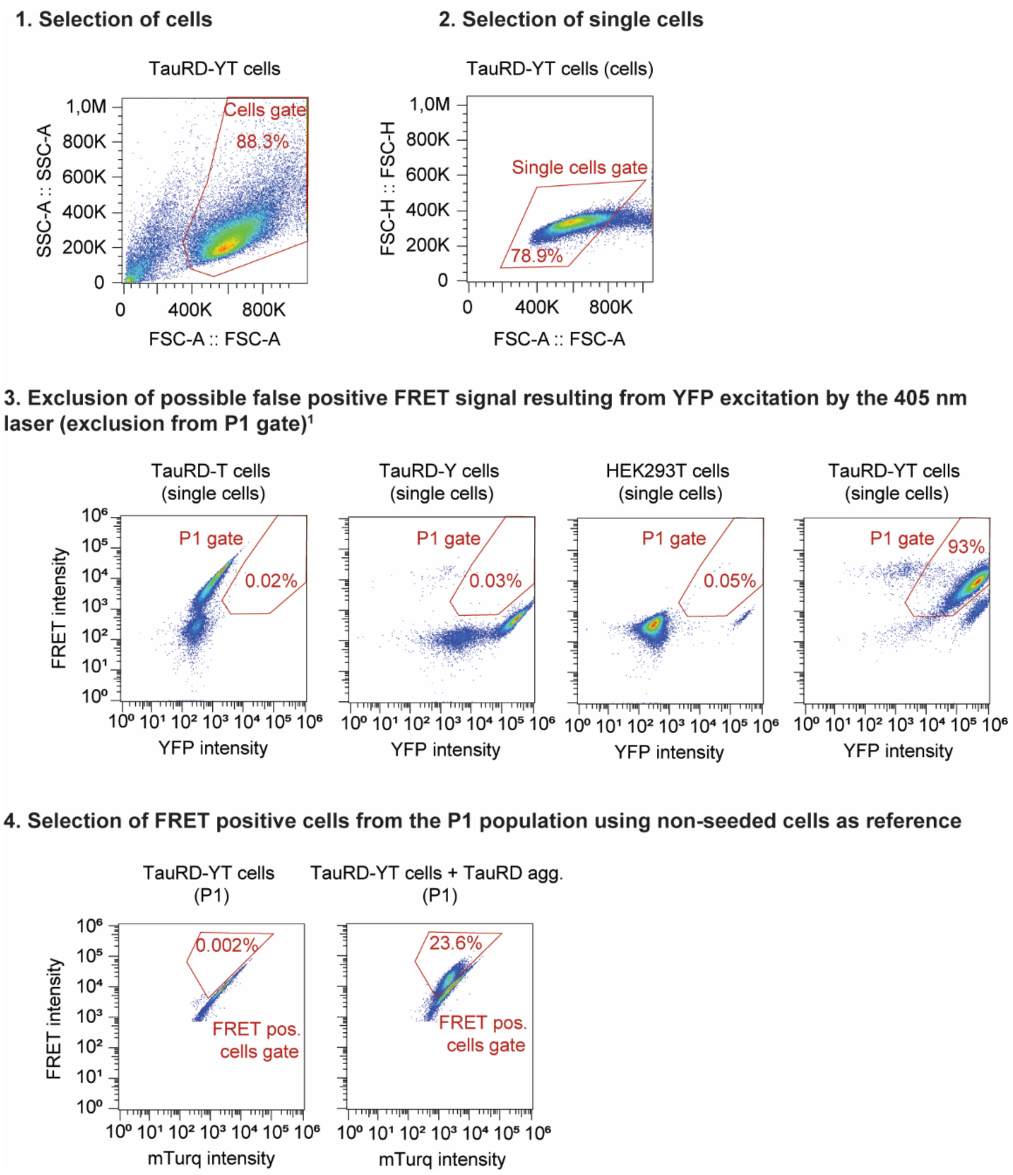


**Supplementary Fig. 10: Flow cytometry strategy for quantification of FRET positive cells.** To measure the mTurquoise2 and FRET fluorescence signals, cells were excited with 405 nm laser light and fluorescence was determined using 440/50 and 530/30 filters, respectively. To measure the YFP fluorescence signal, cells were excited at 488 nm and emission was recorded using a 530/30 filter. For each sample, 50,000 single cells were analyzed. First, cells were selected (1), followed by single cell selection (2). After gating single cells, an additional gate (P1) was introduced to exclude YFP-only cells that show a false-positive signal in the FRET channel due to excitation at 405 nm using as reference TauRD-T cells, TauRD-Y cells and HEK293T cells[1] (3). The FRET positive gate was set by plotting the FRET fluorescence signal versus the mTurquoise2 fluorescence signal using as reference non-seeded cells (4).
